# Immunoglobulins are rapidly internalized by neurons after CNS injury and cleared through lysosomal degradation

**DOI:** 10.64898/2026.07.16.738985

**Authors:** Adrian Castellanos-Molina, Ana Boisvert, Juliette Ferry, Dominic Bélanger, Audrée Laroche, Romain Menet, Frédérique Crépeau, Martine Lessard, Nadia Fortin, Nicolas Vallières, Isabelle Allaeys, Nicolas Bertrand, Ayman ElAli, Éric Boilard, Steve Lacroix

## Abstract

Spinal cord injury (SCI) causes hemorrhage and blood-spinal cord barrier disruption, allowing blood-derived molecules to infiltrate the parenchyma. While immunoglobulins (Ig) are abundant plasma proteins, their distribution and cellular targets within the injured spinal cord remain poorly defined. Here, we show that circulating non-autoimmune immunoglobulins rapidly infiltrate the spinal cord after injury in mice and disseminate beyond the lesion core. IgG, IgM, and IgA accumulate within the parenchyma early post-injury, with IgG displaying the widest spatial distribution, reaching distant spinal segments within hours. Neurons are the predominant cell type internalizing immunoglobulins in the gray matter, whereas astrocytes exhibit moderate uptake in white matter. Intra-cisterna magna administration of fluorescent serum-derived IgG reveals that neurons and astrocytes internalize IgG under physiological conditions, independently of Fc receptor engagement. Although the neonatal Fc receptor (FcRn) has minimal impact on CNS IgG recycling, its genetic deletion significantly improves locomotor recovery after SCI. Both *in vitro* and *in vivo*, neurons clear IgG through lysosomal degradation. Following SCI, inhibition of lysosomal proteases with the clinically approved drug E64d increases CNS IgG retention without compromising locomotor recovery. These findings establish neurons as key targets of circulating immunoglobulins after CNS injury and reveal IgG uptake and clearance pathways that may be leveraged to improve therapeutic performance of monoclonal antibody treatments.

## INTRODUCTION

Spinal cord injury (SCI) affects hundreds of thousands of individuals worldwide each year, most commonly following traumatic events such as vehicle accidents, falls, or sports injuries ^1,2^. The initial physical insult that directly damages the spinal cord is referred to as the primary injury. This initial trauma compromises the structural integrity of the spinal tissue, leading to neuronal and glial cell death, increased permeability of the blood-spinal cord barrier (BSCB), and intraparenchymal hemorrhages. As a result, damage-associated molecular patterns (DAMPs) are released from disrupted cells and blood components infiltrate the spinal cord parenchyma, further altering the local microenvironment ^3^. Several blood-derived molecules are known to play a role in neurodegeneration within the central nervous system (CNS). For instance, fibrinogen and its derivative fibrin can exacerbate pathological processes in conditions such as multiple sclerosis, Alzheimer’s disease, and traumatic CNS injury by interacting with macrophages and microglia, thereby triggering proinflammatory signaling and promoting the initiation and progression of these disorders (reviewed in ^4^).

Among the blood-derived molecules known to abundantly infiltrate sites of CNS injury are immunoglobulins. In circulation, IgG is the most predominant isotype, followed by IgA and IgM. Neurons have been shown to directly interact with IgG. For example, Purkinje cells can internalize IgG in cell culture ^5^. Furthermore, recent studies indicate that self-reactive (auto)antibodies can activate neurons through Fc receptor-dependent signaling, enhancing excitability and contributing to pain ^6,7^. After SCI, both IgG and IgM have been detected within the injured spinal cord tissue during the intermediate and chronic phases ^8,9^. These immunoglobulins are generally thought to be produced locally by B cells activated in response to injury-induced antigens, rather than arising from passive leakage of pre-existing circulating antibodies ^10^. This assumption originated from evidence of elevated titers of autoantibodies against myelin proteins and gangliosides in the serum and cerebrospinal fluid (CSF) of individuals with chronic SCI ^11,12^, as well as the accumulation of B cells at the lesion epicenter in chronically injured mice ^9,10^. Importantly, mice lacking B cells exhibit reduced lesion volume and improved locomotor recovery after SCI ^10^. However, because most research on immunoglobulins in SCI has focused on later post-injury stages, we still know very little about their temporal dynamics, targets, and functional effects in the early stage.

In this study, we aimed to define the spatiotemporal dynamics, cellular targets, and intracellular fate of blood-derived immunoglobulins in the injured CNS. We show that blood-derived immunoglobulins rapidly enter and disseminate throughout the spinal cord after traumatic injury, far beyond the lesion epicenter, where they interact selectively with CNS-resident cells. Notably, neurons and astrocytes intrinsically and predominantly internalize IgG immunoglobulins through an Fc-independent mechanism that is conserved across CNS injury models. Following uptake, immunoglobulins are actively cleared via intracellular degradative pathways, a process that can be modulated without compromising functional recovery after SCI.

## RESULTS

### Immunoglobulins infiltrate and spread beyond the lesion core after contusion SCI

The primary injury associated with the mechanical trauma in SCI compromises vascular integrity, enabling molecules from the periphery to infiltrate the spinal cord parenchyma. Among these components, immunoglobulins, especially IgG, IgM, and IgA, represent some of the most abundant proteins in mammalian blood. To determine whether these immunoglobulins infiltrate the injured spinal cord, we first characterized their spatial and temporal distribution following SCI (Fig. 1A-B). Macroscopically, at 1 day post-injury (dpi), we observed blood surrounding the lesion core, along with a reddish discoloration of the spinal cord extending several millimeters from the lesion epicenter (Fig. 1B). We next performed immunofluorescence staining at multiple timepoints (1 hour, 1 day, and 7 days post-injury) to assess the distribution of IgG, IgM, and IgA within the lesion and adjacent regions (Fig. 1C-N). Immunoglobulins were not detected in the spinal cord parenchyma of uninjured, laminectomized (sham) mice. In contrast, IgG, IgM, and IgA were present over a large portion of the lesion epicenter as early as 1 hour post-injury (hpi). Immunoglobulin abundance peaked at 1 dpi, then declined markedly for IgG and IgM, and reached nearly undetectable levels for IgA by 7 dpi. Quantitative analyses corroborated these observations, demonstrating that the area occupied by immunoglobulin-immunoreactive (IR) staining was maximal between 1 hour and 1 day post-SCI, followed by a significant reduction by Day 7 (Fig. 1O-Q).

**Figure 1.**
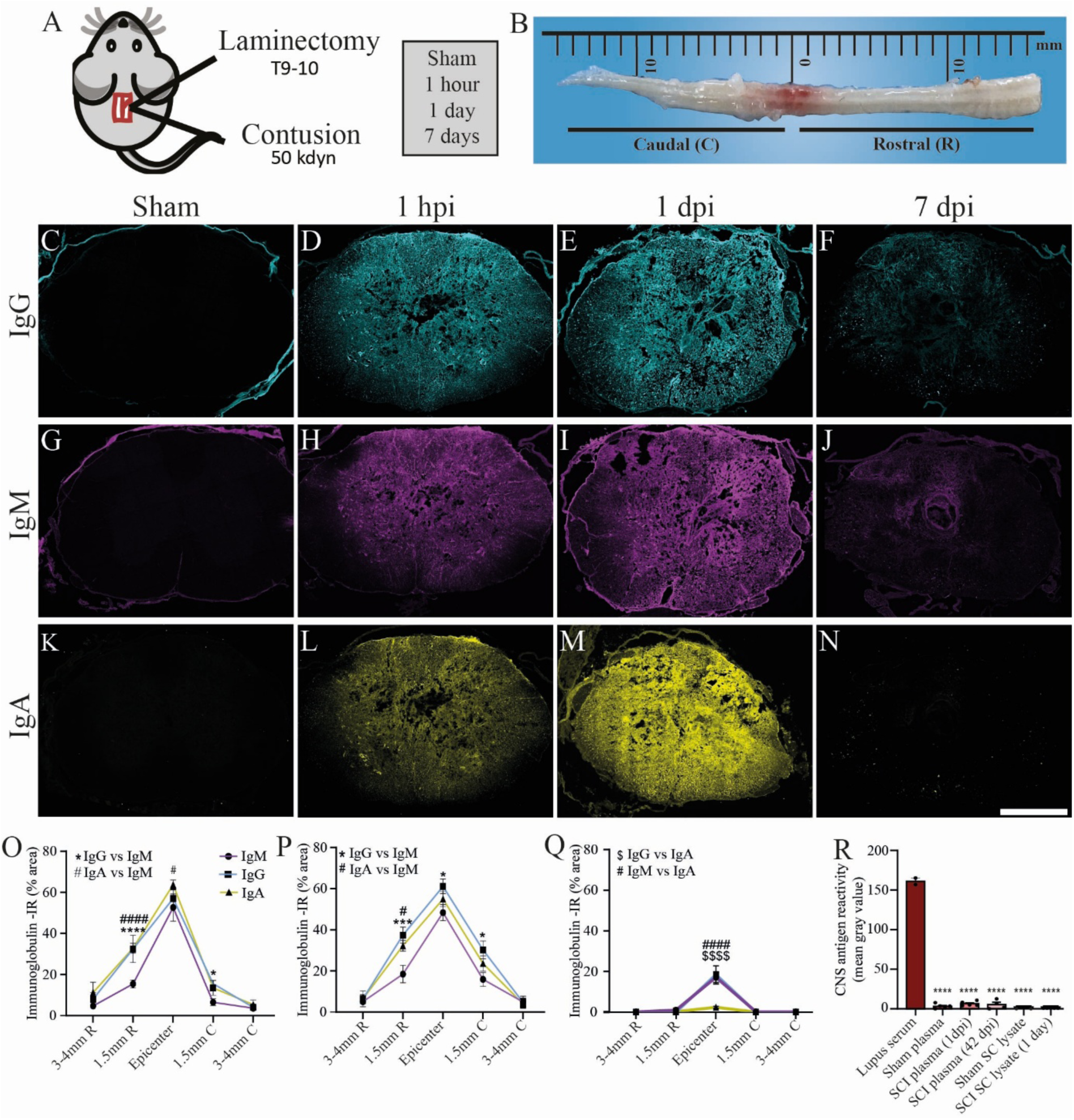
Non-autoimmune immunoglobulins rapidly infiltrate the injured spinal cord and persist through the acute and subacute phases of SCI. (**A**) Schematic representation of the SCI model used. A laminectomy was performed at the T9-10 vertebral level, followed by a 50-kdyn contusion injury. Sham-operated animals underwent laminectomy without contusion. Mice were euthanized at 1 hour, 1 day, or 7 days post-surgery. (**B**) Image of a mouse spinal cord at 1 day post-injury (dpi), showing the lesion site and adjacent rostral (R) and caudal (C) segments. (**C-N**) Representative confocal images showing immunofluorescence for IgG (**C**-**F**, cyan), IgM (**G**-**J**, magenta), and IgA (**K**-**N**, yellow) in coronal spinal cord sections from sham mice (**C**, **G**, **K**) and SCI mice collected at 1 hour (**D**, **H**, **L**), 1 day (**E**, **I**, **M**), and 7 days (**F**, **J**, **N**) post-SCI. (**O-Q**) Quantification of the area occupied by IgG, IgM, and IgA immunostaining at the lesion epicenter and at defined distances rostral (R) and caudal (C) at 1 hour (**O**), 1 day (**P**), and 7 days (**Q**) after SCI. (**R**) Mean pixel intensity in Bio-dot analysis for autoantibody detection using CNS protein as antigen. Serum from lupus-prone NZBWF1 mice was used as a positive control because of its high autoantibody content. Serum samples were collected from sham-operated mice and from SCI mice at 1 or 42 dpi. Protein lysates were prepared from spinal cord segments of sham mice and from the injury site of SCI at 1 dpi. Data are presented as mean ± SEM. Statistical significance was assessed using two-way ANOVA with Bonferroni’s post-hoc test. Pairwise comparisons and corresponding *p*-values are indicated in the graphs, with * p < 0.05, *** p < 0.001, **** p < 0.0001, ^#^ p < 0.05, ^####^ p < 0.0001, and ^$$$$^ p < 0.0001. Scale bar: 250 µm in **C**-**N** (shown in **N**).

Beyond the lesion core, immunoglobulins were also detected at the lesion borders, approximately 1.5 mm from the epicenter, and in more distal regions located 3-4 mm away (Fig. 1O-Q & Suppl. Fig. 1). Notably, at both 1 hour and 1 day post-SCI, the distribution of IgG and IgA, but not IgM, displayed a clear rostro-caudal asymmetry, with greater accumulation rostral to the lesion epicenter (Fig. 1O-P). By 7 dpi, the signal of all three immunoglobulins had markedly diminished throughout the spinal cord (Fig. 1Q). These data indicate that the spinal cord possesses efficient mechanisms to rapidly clear immunoglobulins.

Regarding the spatial distribution of these antibodies, immunoglobulin signal was homogeneously distributed throughout the lesion epicenter (Suppl. Fig. 1). In contrast, at the lesion borders (∼1.5 mm from the epicenter) and in more distal regions (∼3-4 mm), the signal appeared more closely associated with individual cells within both the spinal cord gray and white matter.

We next assessed whether SCI induces the generation of self-reactive autoantibodies. Brain protein lysates were used as antigens in an immunoblot assay and probed with plasma collected from sham mice, as well as plasma from mice sacrificed at 1 and 42 dpi. In addition, spinal cord lysates from sham mice and mice killed at 1 dpi, corresponding to the peak of immunoglobulin presence during acute SCI, were also analyzed (Suppl. Fig. 2A). Serum from lupus-prone NZB/NZW F1 (NZBWF1) mice, which spontaneously develop high titers of circulating autoantibodies, was included as a positive control and, while strong antibody binding was detected in this group, plasma from SCI mice showed no detectable reactivity against CNS proteins and was indistinguishable from sham controls, indicating that SCI does not induce the generation of detectable self-reactive autoantibodies during the acute phase examined (Fig. 1R & Suppl. Fig. 2B). Considering the macroscopic presence of blood and the absence of self-reactive antibodies, it is highly probable that the immunoglobulins observed are predominantly blood-derived rather than newly produced locally.

### Neurons and astrocytes rapidly internalize immunoglobulins after SCI

Immunoglobulins are primarily known for their interactions with immune cells, particularly those expressing Fc receptors (FcR). Given the central role of glial cells, such as microglia and astrocytes, in responding to CNS injury, we sought to determine whether IgG, IgM, and IgA interact with these cell types. At the lesion epicenter, high signal intensity precluded reliable quantification of individual immunoglobulin-expressing cells. We therefore conducted our analyses at the lesion borders, where spinal cord-resident cells were not physically damaged by the injury, or at least did not appear to be.

Microglia express the major Fcγ receptors for IgG, including FcγRI (Cd64), FcγRII (Cd32), and FcγRIII (Cd16), with expression levels strongly upregulated under inflammatory conditions in humans and rodent models ^13-16^. To assess whether microglia interact with immunoglobulins after injury, we used *Cx3cr1*^CreER^::*R26-tdTomato* mice, with tamoxifen induction performed one month prior to SCI as previously described ^17^. This approach ensured selective and persistent tdTomato (tdT) labeling of microglia while minimizing labeling of infiltrating macrophages, which are rapidly replaced due to their high turnover. Surprisingly, microglia exhibited minimal interaction with immunoglobulins (Fig. 2A-F). Although a statistically significant increase in IgG-positive (+) microglia was observed bilaterally 1.5 mm from the lesion epicenter at 1 dpi compared with sham controls, the absolute numbers remained low, reaching only 5.21 ± 2.1 cells/mm² rostrally and 4.86 ± 2.2 cells/mm² caudally (Fig. 2D). Virtually no IgM^+^ and IgA^+^ microglia were detected at these distances.

**Figure 2.**
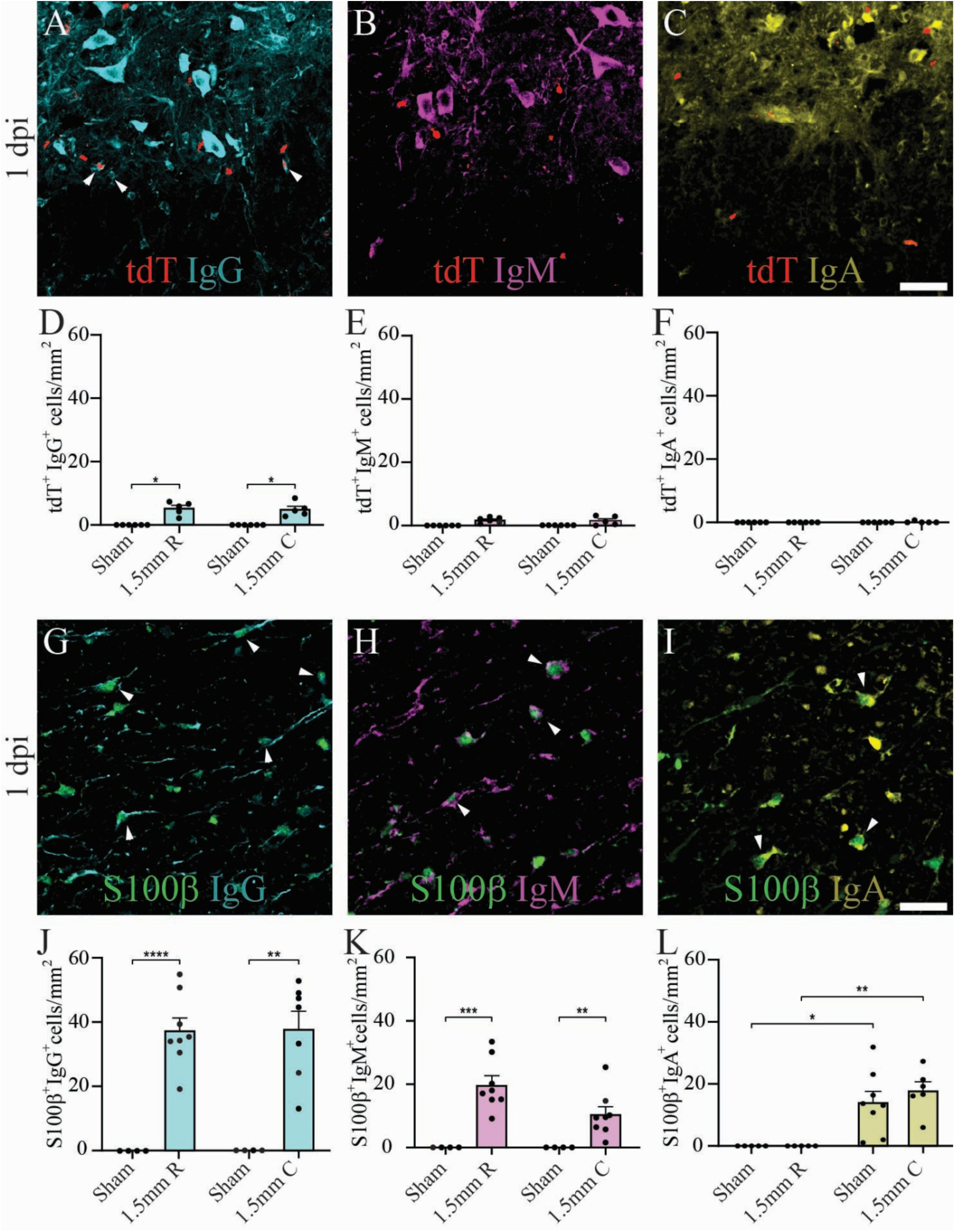
Immunoglobulins that infiltrate the injured spinal cord interact with glial cells. (**A**-**C**) Representative confocal images showing IgG (**A**, cyan), IgM (**B**, magenta), and IgA (**C**, yellow) immunostaining in the spinal cord of injured *Cx3cr1*^Cre^::*Rosa26*^tdT^ mice, in which microglia express the fluorescent reporter tdTomato (tdT, red in **A**-**C**), at 1 day post-injury (dpi). (**D**-**F**) Quantification of tdT⁺ microglia colocalizing with IgG (**D**), IgM (**E**), and IgA (**F**) in sham-operated mice and SCI mice at 1.5 mm rostral (R) and caudal (C) to the lesion epicenter. (**G**-**I**) Representative confocal images showing astrocytes labeled with S100β (green) together with IgG (**G**, cyan), IgM (**H**, magenta), and IgA (**I**, yellow) at 1 dpi. (**J**-**L**) Quantification of S100β⁺ astrocytes colocalizing with IgG (**J**), IgM (**K**), and IgA (**L**). Data are presented as mean ± SEM. Statistical significance was assessed using two-way ANOVA with Bonferroni’s post-hoc test. Pairwise comparisons and corresponding *p*-values are indicated in the graphs, with * p < 0.05, ** p < 0.01, *** p < 0.001, and **** p < 0.0001. Scale bars: 50 µm in **A**-**C** (shown in **C**) and 25 µm **in G**-**I** (in **I**).

We next assessed whether astrocytes interact with immunoglobulins after SCI. The S100β protein is widely used as an astrocytic marker in the CNS, although some studies have reported its expression in subsets of microglia and neurons ^18^. Because our data indicate that microglia exhibit minimal interaction with immunoglobulins, and to avoid potential confounding from S100β^+^ neurons, quantification was restricted to the spinal cord white matter. At 1.5 mm rostral and caudal to the lesion epicenter at 1 dpi, S100β^+^ cells showed clear colocalization with all three immunoglobulins, whereas no such colocalization was detected in the white matter of sham mice (Fig. 2G-L).

Most immunoglobulin-IR cells within the spinal cord gray matter displayed a large neuron-like morphology (Suppl. Fig. 1). To confirm their neuronal identity, we performed colocalization analyses between IgG, IgM, and IgA signals and the pan-neuronal marker NeuN in tissue sections collected from the lesion borders (∼1.5 mm from the epicenter) and from more distal regions (3-4 mm). Confocal immunofluorescence imaging showed that all three immunoglobulins were internalized by neurons, with a significant increase in NeuN^+^ neurons containing IgG, IgM, or IgA at both 1 hour and 1 day post-SCI compared with sham controls (Fig. 3A-O & Suppl. Fig. 3). Quantification revealed that the number of immunoglobulin^+^ neurons was similar at the 1-hour and 1-day time points, but declined markedly by 7 dpi. This neuronal labeling pattern closely mirrored the broader spatial distribution of immunoglobulins observed across time (Fig. 1O-Q). In distal regions located 3-4 mm from the lesion site, we observed a significantly greater number of neurons internalizing immunoglobulins at 1 dpi compared with 1 hpi, likely reflecting the time required for blood-derived immunoglobulins to reach these more distant areas (Fig. 3M-O). Overall, these findings demonstrate that neurons are major targets for immunoglobulin uptake after SCI.

**Figure 3.**
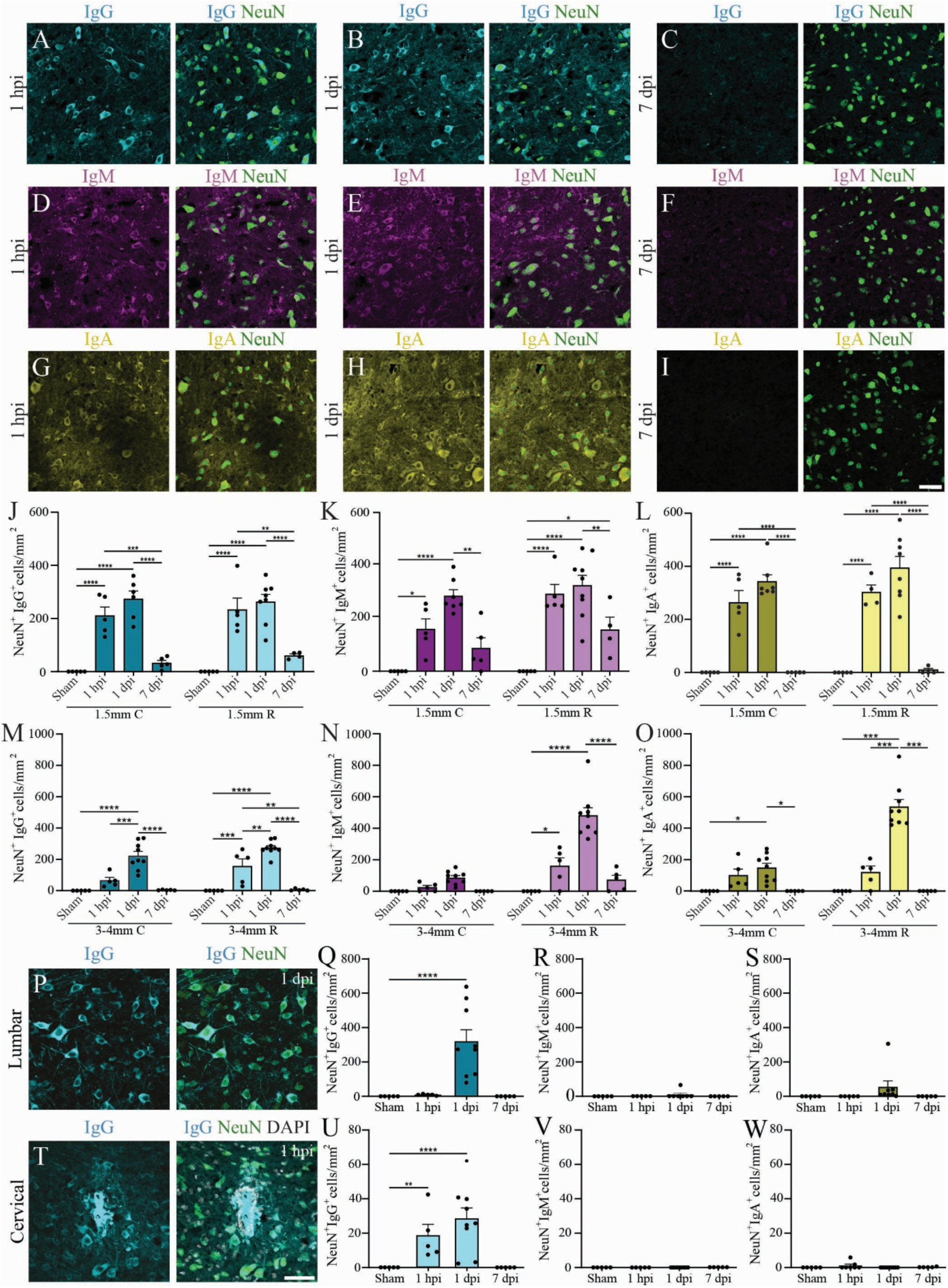
Neurons internalize immunoglobulins after SCI. (**A**-**I**) Representative confocal images showing neurons labeled with NeuN (green) together with IgG (**A**-**C**, cyan), IgM (**D**-**F**, magenta), and IgA (**G**-**I**, yellow) at 1 hour (**A**, **D**, **G**), 1 day (**B**, **E**, **H**), and 7 days (**C**, **F**, **I**) after SCI, imaged 1.5 mm rostral to the lesion epicenter. (**J**-**O**) Quantification of NeuN^+^ neurons colocalizing with IgG (**J**, **M**), IgM (**K**, **N**), and IgA (**L**, **O**) in sham-operated mice and in SCI mice at 1.5 mm (**J**-**L**) and 3-4 mm (**M**-**O**) rostral (R) and caudal (C) to the lesion epicenter. Mice were euthanized at 1 hour post-injury (hpi), 1 day post-injury (dpi), or 7 dpi. (**P**) Representative confocal images showing NeuN^+^ IgG^+^ neurons in a lumbar spinal cord coronal section at 1 dpi. (**Q**-**S**) Quantification of NeuN^+^ IgG^+^ (**Q**), NeuN^+^ IgM^+^ (**R**), and NeuN^+^ IgA^+^ (**S**) neurons in the lumbar spinal cord of sham-operated mice at 1 day post-surgery, and of SCI mice at 1 hpi, 1 dpi, and 7 dpi. (**T**) Confocal images showing NeuN^+^ IgG^+^ neurons in the cervical spinal cord at 1 hpi. (**U**-**W**) Quantification of NeuN^+^ IgG^+^ (**U**), NeuN^+^ IgM^+^ (**V**), and NeuN^+^ IgA^+^ (**W**) neurons in the cervical spinal cord of sham-operated mice at 1 day post-surgery and SCI mice at 1 hpi, 1 dpi, and 7 dpi. Data are presented as mean ± SEM. Statistical significance was assessed using two-way ANOVA with Bonferroni’s post-hoc test. Pairwise comparisons and corresponding *p*-values are indicated in the graphs, with * p < 0.05, ** p < 0.01, *** p < 0.001, and **** p < 0.0001. Scale bars: 50 µm in **A**-**I** (shown in **I**) and 50 µm in both **P** and **T** (in **T**).

To determine whether immunoglobulin infiltration extended beyond the thoracic region, we examined more distal spinal segments, specifically the L5-L6 lumbar and C3-C4 cervical levels. In the lumbar spinal cord, IgG was detected only at 1 dpi, whereas IgM and IgA signals were minimal or absent (Fig. 3P-S). Strikingly, IgG signal was present in the cervical spinal cord as early as 1 hpi. In this region, IgG^+^ neurons were primarily localized around the central canal, and IgG accumulation was also observed in the extracellular space (Fig. 3T-U). IgG remained detectable in the C3-C4 region at 1 dpi, but was no longer observed at 7 dpi (Fig. 3U). In contrast, neither IgM nor IgA was detected in the cervical spinal cord at any time point examined (Fig. 3V-W).

Notably, NeuN immunostaining revealed a progressive reduction in neuronal density at the lesion epicenter at 1 hour, 1 day, and 7 days post-SCI compared with sham animals (Suppl. Fig. 4A-D). In contrast, regions located 1.5 mm rostral and caudal to the lesion site displayed a preserved neuronal distribution across all time points examined, with no significant changes in neuronal numbers over time (Suppl. Fig. 4F-M). Given the large proportion of neurons that internalize immunoglobulins in these regions, these data suggest that immunoglobulin internalization does not promote neuronal loss.

Altogether, these findings indicate that immunoglobulins preferentially associate with astrocytes in the spinal cord white matter and with neurons in the gray matter. Consistent with this observation, a substantial proportion of neurons in regions adjacent to the lesion site rapidly internalize immunoglobulins without any apparent impact on neuronal survival, although this conclusion warrants more rigorous investigation. Moreover, our results suggest that IgG may exploit the central canal as a conduit to rapidly travel long distances and penetrate into the CNS once the blood-CSF barrier becomes compromised, such as after SCI.

### Immunoglobulin internalization and neuronal uptake exhibit similar temporal dynamics in a mouse model of ischemic injury

Ischemic stroke acutely disrupts the blood-brain barrier (BBB) by compromising tight junction integrity and increasing endothelial permeability ^19,20^. This disruption facilitates the leakage of plasma proteins and the infiltration of immune cells into the brain parenchyma, thereby exacerbating edema, inflammation, and neuronal injury. To determine whether neurons outside the spinal cord also internalize immunoglobulins following a CNS insult, we used the middle cerebral artery occlusion (MCAO) mouse model, which induces transient ischemia and neuronal loss in the cortex and striatum (Suppl. Fig. 5A). We assessed immunoglobulin internalization throughout the affected region by examining coronal brain sections at both the striatal and hippocampal levels (Suppl. Fig. 5B). The lesion area was defined as the region in which NeuN signal intensity was reduced on the ipsilateral side compared to the contralateral side (Suppl. Fig. 5C-D). The presence of IgG, IgM, and IgA was indicated by a marked increase in immunofluorescence intensity, strictly localized within the lesion borders (Suppl. Fig. 5E-P). Notably, the different immunoglobulin classes exhibited distinct post-MCAO infiltration profiles. IgG was detected as early as 1 day after MCAO, covering nearly the entire lesion and indicating a rapid BBB disruption (Supp. Fig. 5E-F). IgG remained detectable at 7 days in both the hippocampal and striatal sections (Supp. Fig. 5G-H). In contrast, almost no IgM was present in the lesion at 1 day (Supp. Fig. 5I-J), but IgM accumulation markedly increased by 7 days post-MCAO (Supp. Fig. 5K-L). This delayed entry likely reflects both its larger molecular size and its lower serum abundance relative to IgG, factors that may limit its diffusion across a partially disrupted BBB. IgA followed a similar delayed pattern, being undetectable at 1 day but clearly present within the lesion by 7 days post-MCAO (Suppl. Fig. 5M-P).

**Figure 4.**
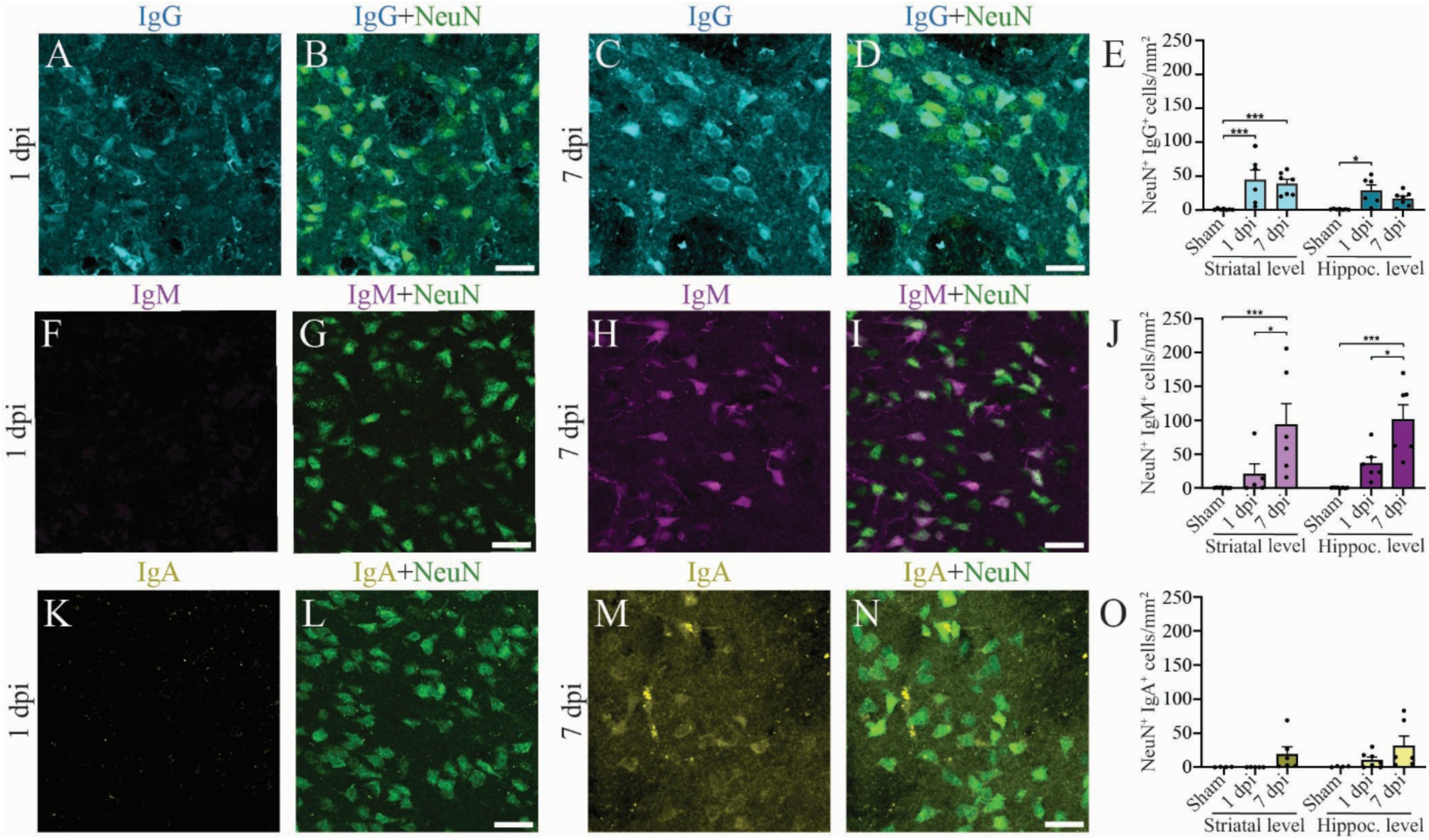
Neurons internalize immunoglobulins after stroke. (**A**-**D**) Representative confocal images showing IgG (blue) internalized within NeuN^+^ neurons (green) in the ischemic core at the striatal level at 1 day post-ischemia (dpi, **A**-**B**) and 7 dpi (**C**-**D**) following middle cerebral artery occlusion (MCAO) in mice. (**E**) Quantification of NeuN^+^ IgG^+^ neurons within the ischemic lesion in coronal brain sections taken at the striatal and hippocampal levels in sham-operated mice (Day 1) and MCAO mice at 1 and 7 dpi. (**F**-**I**) Confocal images showing IgM (magenta) internalized within NeuN^+^ neurons (green) in the ischemic lesion at 1 dpi (**F**-**G**) and 7 dpi (**H**-**I**) at the striatal level in MCAO mice. (**J**) Quantification of NeuN^+^ IgM^+^ neurons at the striatal and hippocampal levels in sham-operated (Day 1) and MCAO mice at 1 and 7 dpi. (**K**-**N**) Confocal images showing IgA (yellow) internalized within NeuN^+^ neurons (green) in the ischemic lesion at 1 dpi (**F**-**G**) and 7 dpi (**H**-**I**) at the striatal level in MCAO mice. (**O**) Quantification of NeuN^+^ IgA^+^ neurons at the striatal and hippocampal levels in sham-operated (Day 1) and MCAO mice at 1 and 7 dpi. Data are presented as mean ± SEM. Statistical significance was assessed using two-way ANOVA with Bonferroni’s post-hoc test. Pairwise comparisons and corresponding *p*-values are indicated in the graphs, with * p < 0.05 and *** p < 0.001. Scale bars: 25 µm in **A**-**D** (shown in **B** and **D**), 25 µm in **F**-**I** (in **G** and **I**), and 25 µm in **K**-**N** (in **L** and **N**).

**Figure 5.**
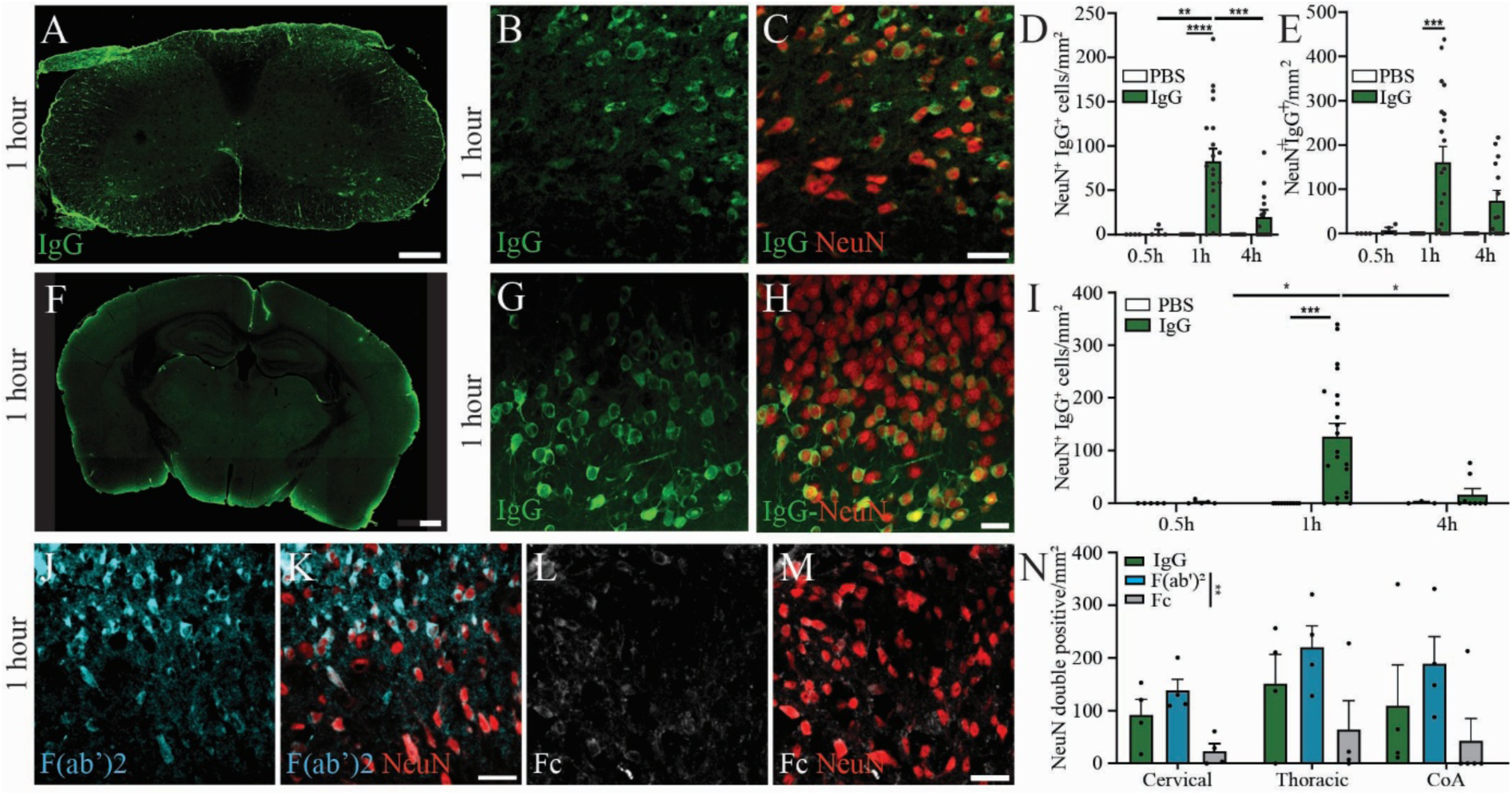
Neuronal IgG uptake occurs independently of injury and is mediated primarily by the F(ab’)_2_ domain rather than the Fc domain. (**A**-**C**) Representative confocal images showing the distribution of Alexa488-conjugated IgG (green) in the cervical spinal cord at 1 hour following intra-cisterna magna (i.c.m.) injection (**A**), and its internalization within NeuN^+^ neurons (red) in higher-magnification views (**B**-**C**). (**D**-**E**) Quantification of NeuN^+^ IgG^+^ neurons in the cervical (**D**) and thoracic (**E**) spinal cord at 0.5, 1, and 4 hours (h) after injection of Alexa488-conjugated IgG or PBS. (**F**-**H**) Representative confocal images showing the distribution of Alexa488-conjugated IgG (green) in the brain at the hippocampal level at 1h post-i.c.m. injection (**F**), and its internalization within NeuN^+^ neurons (red) in higher-magnification views in the cortical amygdala area (CoA)(**G**-**H**). (**I**) Quantification of NeuN^+^ IgG^+^ neurons in the CoA at 0.5, 1, and 4 hours after injection of either Alexa488-conjugated IgG or PBS. (**J**-**K**) Representative confocal images showing the internalization of F(ab’)₂ fragments (cyan) within NeuN^+^ neurons (red) in the spinal cord at 1h post-i.c.m. injection. (**L**-**M**) Representative confocal images showing the minimal internalization of Fc fragments (grey) within NeuN^+^ neurons (red) of the spinal cord. (**N**) Quantification of NeuN⁺ neurons colocalizing with whole IgG, F(ab’)₂ fragments, or Fc fragments in the cervical and thoracic spinal cord and in the CoA of the brain at 1h post-i.c.m. injection. Data are presented as mean ± SEM. Statistical significance was assessed using two-way ANOVA with Bonferroni’s post-hoc test. Pairwise comparisons and corresponding *p*-values are indicated in the graphs, with * p < 0.05, ** p < 0.01, *** p < 0.001, and **** p < 0.0001. Scale bars: 250 µm in **A**, 25 µm in **B**-**C** (shown in **C**), 500 µm in **F**, 25 µm in **G**-**H** (in **H**), and 25 µm in **J**-**M** (in **K** and **M**).

Similar to our observations in SCI, neurons also exhibited the capacity to internalize IgG following MCAO (Fig. 4A-E). Neurons that survived within the ischemic territory displayed an atrophic morphology with reduced NeuN signal intensity (Suppl. Fig. 5C-D), yet internalized IgG as early as at 1 day post-ischemia (dpi) in both the striatum and hippocampus (Fig. 4A-B). Unlike SCI, where immunoglobulin presence is nearly absent by 7 dpi (Fig. 1), IgG continued to colocalize with neurons at 7 days after MCAO, showing only a small, non-significant reduction compared with Day 1 (Fig. 4A-E). Neuronal internalization of IgM was minimal at Day 1, with virtually no IgM^+^ neurons detected in the striatum (Fig. 4F-G). However, a robust increase in IgM colocalization with NeuN was observed by Day 7 (Fig. 4H-J). In contrast to SCI, where IgA internalization occurs at levels comparable to other immunoglobulins (Fig. 3), only a small proportion of neurons internalized IgA after MCAO, both at 1 and 7 dpi and in both the striatal and hippocampal regions (Fig. 4K-O).

Overall, these findings demonstrate that ischemic injury leads to sustained immunoglobulin accumulation within the ischemia core, with neurons showing robust early IgG uptake and a marked increase in IgM internalization by Day 7. Together with our SCI data, these results suggest that neuronal antibody internalization may represent a common acute response to CNS injury.

### Neurons, astrocytes and blood vessels internalize IgG constitutively under homeostatic conditions

Because IgG is the predominant immunoglobulin in normal mouse serum and is consistently detected across CNS injury models, and given that therapeutic monoclonal antibodies are almost exclusively IgG isotypes, we focused our mechanistic analyses on IgG. To determine whether CNS-resident cells can internalize serum IgG under physiological, non-injured conditions, we injected Alexa Fluor 488-conjugated IgG purified from naïve mouse serum into the cisterna magna (i.c.m.), a route that provides direct access to CNS parenchyma. We then examined cervical and thoracic spinal cord sections, as well as the cortical amygdala (CoA), a region known to receive cisterna magna-delivered substances ^21^. This approach allowed us to assess whether simple exposure to circulating IgG, in the absence of injury-induced stress or apoptosis, is sufficient to trigger its uptake by CNS cells. Thirty minutes after injection, IgG signal was largely confined to the meninges in both the spinal cord and brain, with minimal internalization within the CNS parenchyma (Suppl. Fig. 6A-D). By 1 hour post-injection, IgG had penetrated the spinal cord parenchyma, including both white and gray matter (Fig. 5A & Suppl. Fig. 6E-F), and was readily internalized by neurons at both cervical and thoracic levels (Fig. 5B-E). A similar pattern was observed in the brain, where neuronal IgG uptake peaked at 1 hour in the CoA (Fig. 5F-I & Suppl. Fig. 6G-H). Four hours after injection, overall IgG signal in both the spinal cord and brain was markedly reduced in most mice (Suppl. Fig. 6I-L), suggesting rapid clearance or degradation. Neurons also appeared to clear internalized IgG, as indicated by a significant reduction in the density of IgG-colocalizing neurons at 4 hours compared with the 1-hour time point (Fig. 5D-E & I).

Given the presence of IgG within the white matter, we next investigated whether glial cells and blood vessels also internalize IgG under normal physiological conditions. Several blood vessels located near the meningeal surface in both the spinal cord and brain compartments appeared IgG^+^ 30 minutes after i.c.m. injection (Suppl. Fig. 7A-H). In addition, we performed immunostaining for the astrocytic marker S100β and the microglial marker Iba1 to assess colocalization with IgG (Suppl. Fig. 7I-O). Consistent with our observations in SCI, IgG showed minimal colocalization with Iba1^+^ microglia, whereas nearly half of S100β^+^ astrocytes displayed robust IgG uptake. These findings indicate that astrocytes actively internalize IgG even in the absence of injury. The fact that both astrocytes and neurons internalize IgG under baseline conditions, and after SCI and stroke, suggests an intrinsic capacity of these CNS-resident cells to interact with immunoglobulins, a mechanism that may be relevant across CNS pathologies characterized by IgG infiltration.

### Neurons constitutively internalize IgG through an Fc-independent mechanism

In principle, IgG uptake by murine cells occurs via two non-exclusive routes: (i) Fc-dependent engagement of low and high-affinity Fcγ receptors (e.g., Cd64, Cd32b, Cd16) that trigger endocytosis or phagocytosis of IgG-immune complexes or monomeric IgG, and (ii) Fab-dependent binding of IgG to cell-surface antigens followed by receptor-mediated internalization of the antibody-antigen complex ^22^. To further explore the mechanism underlying neuronal IgG uptake, we examined whether internalization depends on Fc-mediated and/or Fab-mediated recognition. To achieve this, Alexa Fluor 488-conjugated Fc fragments and F(ab’)_2_ fragments of IgG were injected into the cisterna magna (Fig. 5J-N). Surprisingly, neurons robustly internalized the F(ab’)_2_ fragment, which lacks the Fc region required for Fc receptor binding, at levels comparable to, or slightly exceeding, those observed with full-length IgG. By contrast, although not entirely absent, uptake of the Fc fragment alone was significantly reduced relative to the F(ab’)_2_ fragment (Fig. 5L-N), suggesting that the Fc region may be unnecessary for IgG internalization under non-injured conditions.

Altogether, these findings suggest that IgG uptake does not rely on Fc receptor engagement but instead depends on the F(ab’)_2_ portion, pointing to an intrinsic, physiological mechanism that likely reflects a fundamental feature of CNS cell biology.

### Lysosomes drive neuronal degradation of internalized IgG

The neonatal Fc receptor (FcRn) is known to protect IgG from lysosomal degradation by promoting its recycling back to the cell surface, thus extending its half-life^23^. However, in FcRn^+^cells, IgG that fails to bind FcRn is directed to lysosomes for degradation^24^. Given the progressive decline in IgG levels within the CNS parenchyma (Figs. 3 & 5), we hypothesized that neurons may actively degrade IgG via lysosomal pathways and/or an FcRn-dependent mechanism.

To investigate the mechanisms of IgG elimination, we isolated cortical neurons from P0 *Emx1*^Cre^::*Rosa26*^tdT^ pups, whose neurons endogenously express the fluorescent protein tdT. After 7 days of *in vitro* maturation, the neurons were incubated with IgG for 48 hours, confirming their ability to internalize IgG (see Video 1). Following incubation, IgG was removed from the culture medium, and the neurons were maintained in IgG-free medium for 1, 4, or 24 hours to assess their capacity to degrade the internalized IgG (Fig. 6A). Immunofluorescence imaging showed a progressive decline in IgG signal intensity over the 24-hour period following IgG removal (Fig. 6B-H). We next quantified the relative IgG signal intensity compared with the negative control. After 48 hours of IgG incubation, the signal was approximately twice that of the negative control (Fig. 6H). By 4 hours after IgG removal, a significant decrease was evident, with values dropping to ∼1.6 and remaining at a similar level at 24 hours. These results indicate that neurons may actively clear internalized IgG *in vitro*.

**Figure 6.**
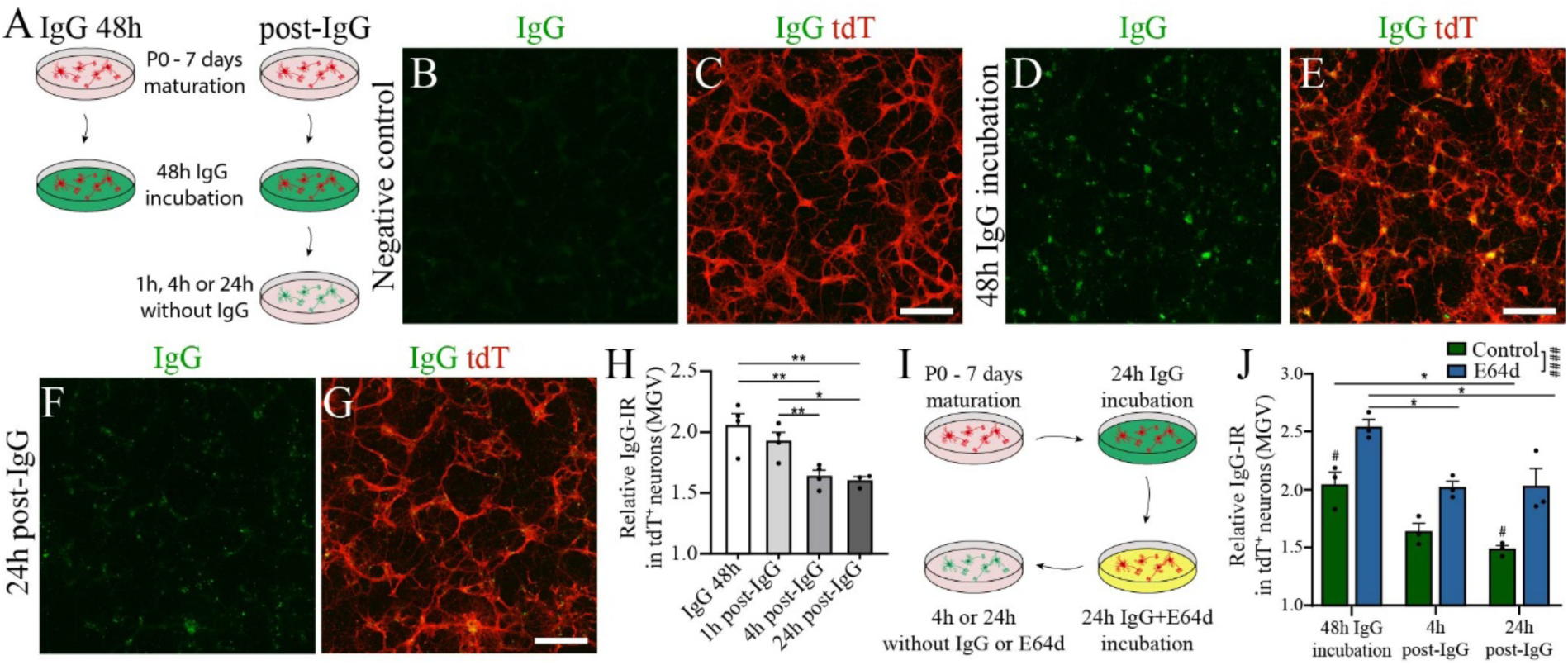
Neurons degrade IgG via the endolysosomal pathway. (**A**) Schematic illustrating the experimental design used to examine IgG internalization in primary mouse cortical neurons. (**B**-**E**) Representative immunofluorescence images showing no detectable IgG internalization (green) in tdT^+^ cortical neurons (red) from P0 *Emx1*^Cre^::*Rosa26*^tdT^ pups in the negative control (no IgG, **B**-**C**), and robust IgG uptake after 48 hours of IgG incubation (**D**-**E**). (**F**-**G**) Representative immunofluorescence image showing neuronal IgG degradation 24 hours after removal of IgG from the culture medium, following a 48-hour internalization period. (**H**) Quantification of the average intensity of IgG signal colocalizing with tdT, normalized to the negative control, after 48 hours of IgG preincubation and subsequent IgG withdrawal for 1, 4, or 24 hours. (**I**) Schematic illustrating the experimental design used to assess how E64d-mediated inhibition of the endolysosomal pathway affects IgG degradation in primary cortical neurons. (**J**) Quantification of tdT-colocalized IgG signal intensity, normalized to the negative control, after 48 hours of IgG preincubation and subsequent IgG withdrawal for 1, 4, or 24 hours in the presence or absence of E64d. Data are presented as mean ± SEM. Statistical significance was assessed using two-way ANOVA with Bonferroni’s post-hoc test. Pairwise comparisons and corresponding *p*-values are indicated in the graphs, with * p < 0.05, ** p < 0.01, ^#^ p < 0.05, and ^####^ p < 0.0001. Scale bar: 250 µm in **B-G** (shown in **C**, **E**, and **G**).

To determine whether this degradation is mediated by the lysosome, we repeated the experiment described above but added E64d, an inhibitor of cysteine proteases, including lysosomal cathepsins ^2,25^, thus disrupting lysosomal function during the final 24 hours of IgG incubation (Fig. 6I). Neurons treated with E64d showed a significant increase in IgG signal intensity compared with control cultures (Fig. 6J). However, lysosomal inhibition did not fully prevent the decline in IgG intensity after IgG removal from the medium, perhaps because lysosomal inhibition was incomplete, or because additional mechanisms may also be involved. Together, these results indicate that lysosomal activity contributes to neuronal IgG clearance.

### Inhibition of lysosomal activity increases IgG retention in the spinal cord and is well tolerated after SCI

We next asked whether IgG undergoes endolysosomal degradation in neurons *in vivo*, and if so, how inhibition of this pathway influences IgG handling after SCI. Because E64d has been reported to cross the BBB ^26^, we administered it intraperitoneally (i.p.) 1 hour before i.c.m. injection of IgG-A488, then quantified by immunofluorescence the number of neurons that internalized IgG in the cervical, thoracic, and lumbar spinal cord gray matter. One hour after IgG injection, E64d-treated mice showed a modest increase in NeuN^+^ IgG^+^ neurons in the cervical and thoracic cord compared with vehicle controls, although these differences did not reach statistical significance (Fig. 7A). A similar, non-significant trend was observed in the cervical white matter, where E64d-treated mice exhibited slightly increased IgG signal relative to controls (Fig. 7B). IgG levels in the thoracic and lumbar white matter were comparable between vehicle- and E64d-treated mice. Notably, the situation changed at 4 hours post-injection, as the number of NeuN^+^ IgG^+^ neurons increased at all spinal cord levels in E64d-treated mice compared with vehicle-treated controls, indicating enhanced neuronal IgG uptake in response to lysosomal inhibition (Fig. 7C). On average, vehicle-treated mice displayed 66.7 ± 30.8, 131.2 ± 46.7, and 89.8 ± 32.1 neurons/mm² in the cervical, thoracic, and lumbar cord, respectively, whereas E64d-treated mice showed higher counts of 134.2 ± 37.8, 340.5 ± 79.4, and 199.32 ± 34.2 neurons/mm². Although individual spinal segments did not reach statistical significance when analyzed separately, which likely reflects the intrinsic variability in the model, combined analysis across the three spinal cord segments revealed a significant increase in neuronal IgG internalization in E64d-treated mice at 4 hours post-injection. This increase paralleled a rise in the area occupied by IgG signal, which was significantly greater in the thoracic cord of E64d-treated animals compared with vehicle-treated controls (Fig. 7D).

**Figure 7.**
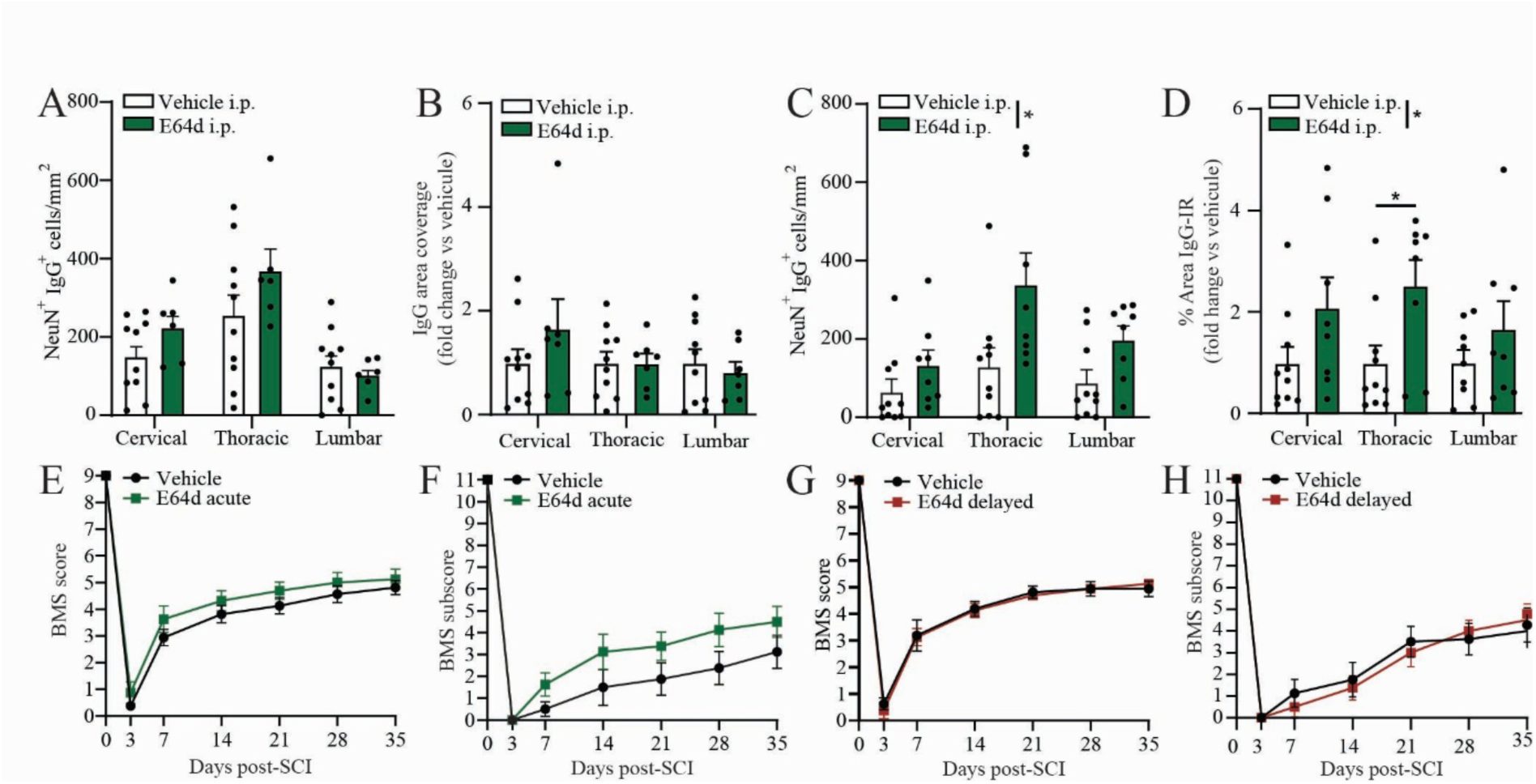
Inhibition of lysosomal activity with E64d in mice reduces neuronal IgG degradation following i.c.m. IgG injection. (**A**) Quantification of NeuN⁺ IgG⁺ neurons in the cervical, thoracic, and lumbar spinal cord gray matter of E64d-pretreated and vehicle-pretreated mice 1 hour after i.c.m. IgG injection. (**B**) Quantification of the area occupied by IgG signal in the spinal cord white matter of E64d-pretreated and vehicle-pretreated mice 1 hour after i.c.m. IgG injection. Data are presented as fold change relative to vehicle-treated controls. (**C**) Quantification of NeuN⁺ IgG⁺ neurons in the spinal cord gray matter of E64d-pretreated and vehicle-pretreated mice 4 hours after IgG injection. (**D**) Quantification of the area occupied by IgG signal in the spinal cord white matter of E64d-pretreated and vehicle-pretreated mice 4 hours after IgG injection. (**E**-**H**) Locomotor function was assessed using the BMS score (**E**, **G**) and BMS subscore (**F**, **H**) in SCI mice treated with E64d or vehicle 1 day (acute treatment) or 14 days (delayed treatment) after injury (n = 8 mice/group). Data are presented as mean ± SEM. Statistical significance was determined by a two-way repeated measures ANOVA with Bonferroni’s post-hoc test. Pairwise comparisons and corresponding *p*-values are indicated in the graphs, with * p < 0.05.

Because lysosomal inhibition with E64d increased IgG retention in the spinal cord, blocking IgG degradation may represent a viable combinatorial strategy to enhance the stability and tissue persistence of monoclonal antibody-based therapies in the context of SCI. However, determining the suitability of E64d as an adjuvant requires confirming that it does not exacerbate post-injury pathology. To address this, E64d was administered intraperitoneally either acutely, at 1 day post-SCI or in a delayed paradigm at 14 days post-SCI, and locomotor recovery was evaluated longitudinally using the Basso Mouse Scale (BMS). Acute E64d administration produced a modest trend toward improved locomotor recovery compared with vehicle-treated mice, a tendency that was more apparent in the BMS tally subscore, although it did not reach statistical significance (Fig. 7E-F). In contrast, delayed E64d administration at 14 days post-injury neither improved nor impaired locomotor recovery (Fig. 7G-H). Overall, these results indicate that E64d does not exert detrimental effects on functional recovery after SCI and support its potential use as a safe adjuvant strategy to improve the persistence and therapeutic efficacy of monoclonal antibodies within the injured spinal cord.

### FcRn deficiency does not affect spinal IgG uptake and retention yet significantly improves locomotor recovery after SCI

In peripheral tissues, FcRn is a key regulator of IgG recycling and stability ^23^. We next examined whether FcRn deficiency alters neuronal IgG uptake or retention using FcRn-knockout (KO) mice injected i.c.m. with IgG-A488 and sacrificed 1 or 4 hours after injection. At 1 hour post-injection, FcRn-KO mice showed no significant differences in the number of NeuN^+^ IgG^+^ neurons across cervical, thoracic, or lumbar spinal cord segments compared with WT controls (Fig. 8A). Consistently, the area occupied by IgG signal in the spinal cord white matter did not differ between genotypes at this early time point (Fig. 8B). These data indicate that initial IgG uptake by neurons and astrocytes is unaffected by the absence of FcRn. Similarly, at 4 hours post-injection, FcRn-KO mice showed no significant differences in the number of NeuN^+^ IgG^+^ neurons across all three spinal cord segments (Fig. 8C), or in IgG accumulation throughout the spinal cord white matter (Fig. 8D). However, although the NeuN^+^ IgG^+^ neurons did not reach statistical significance, it showed a trend toward elevation in FcRn-KO mice compared with WT animals at 4 hours post-injection.

**Figure 8.**
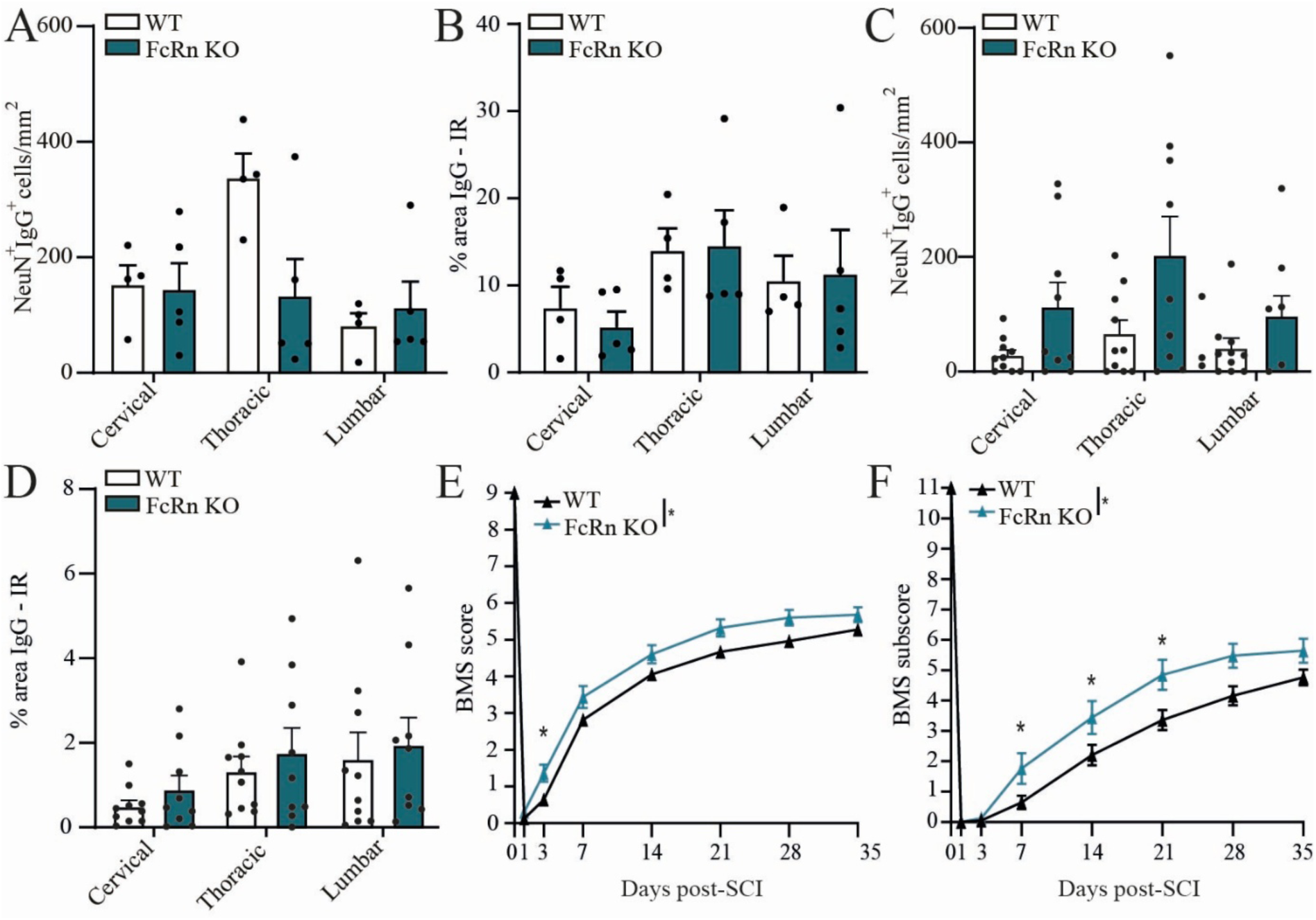
FcRn deficiency improves locomotor recovery after SCI independently of spinal IgG uptake and retention. (**A**) Quantification of NeuN⁺ IgG⁺ neurons in the cervical, thoracic, and lumbar spinal cord gray matter of FcRn-KO and wild-type (WT) mice 1 hour after i.c.m. IgG injection. (**B**) Quantification of the area occupied by IgG signal in the spinal cord white matter of FcRn-KO and WT mice 1 hour after IgG administration. (**C**) Quantification of NeuN⁺ IgG⁺ neurons in the spinal cord of FcRn-KO and WT mice 4 hours after IgG injection. (**D**) Quantification of the area occupied by IgG signal in the spinal cord white matter of FcRn-KO and WT mice 4 hours after IgG injection. (**E**-**F**) Locomotor function was assessed using the BMS score (**E**) and BMS subscore (**F**) after SCI in FcRn-KO (n = 25 mice) and WT mice (n = 40 mice). Data are presented as mean ± SEM. Statistical significance was determined by a two-way repeated measures ANOVA with Bonferroni’s post-hoc test. Pairwise comparisons and corresponding *p*-values are indicated in the graphs, with * p < 0.05.

Since FcRn has been shown to mediate IgG transport across biological barriers ^27,28^, and is expressed at the BBB ^29^, we next investigated whether IgG injected into the CSF could still be transported into the circulation in the absence of this receptor. Because rabbit IgG exhibits high affinity for FcRn and can be readily distinguished from endogenous murine IgG ^30^, it was injected i.c.m. and quantified in plasma 4 hours later (Suppl. Fig. 8A). While rabbit IgG was undetectable in the blood of naïve mice, a substantial fraction was present in the plasma following i.c.m. injection, with FcRn-KO and WT mice having comparable levels of rabbit IgG in plasma 4 hours post-injection (Suppl. Fig. 8B). To determine whether other Fcγ receptors contribute to antibody efflux from the CSF and CNS parenchyma into the circulation, we performed the same experiment in mice lacking the activating FcγRI, FcγRIII, and FcγRIV receptors (FcγR-KO) and in mice lacking the inhibitory receptor FcγRIIb (FcγRIIb-KO). Similar to FcRn-KO mice, neither FcγR-KO nor FcγRIIb-KO mice exhibited altered plasma levels of rabbit IgG compared with their respective WT controls (Suppl. Fig. 8C-D). Together, these findings indicate that IgG is rapidly transferred from the CSF and CNS parenchyma to the bloodstream through an Fc receptor-independent mechanism.

We next examined whether FcRn deficiency influences locomotor recovery after SCI. As shown in Fig. 8E, FcRn-KO mice exhibited significantly higher BMS scores than WT mice as early as 3 days post-injury. Analysis of BMS subscores further revealed enhanced fine motor coordination and stepping performance in FcRn-KO mice at 7, 14 and 21 days post-SCI (Fig. 8F). Collectively, these findings suggest that, despite having no apparent role in neuronal IgG uptake, retention, or trafficking, FcRn may represent a promising therapeutic target for improving functional outcomes after SCI.

## DISCUSSION

This study shows that immunoglobulins rapidly infiltrate the injured spinal cord, spread far beyond the lesion core, and are robustly internalized by CNS-resident cells, particularly neurons. IgG, IgM, and IgA that do not appear to be self-reactive antibodies enter the contused spinal cord within 1 hour, covering large areas of the lesion and extending several millimeters into surrounding tissue, revealing an unexpectedly broad capacity of peripheral antibodies to disseminate through the injured CNS. Across both SCI and ischemic stroke models, neurons consistently displayed the highest immunoglobulin uptake, with astrocytes showing moderate and microglia minimal internalization. Through i.c.m. injection of fluorescent IgG, we further demonstrate that neurons and astrocytes internalize serum IgG even in the absence of CNS injury, independently of Fc-receptor engagement. Once internalized, neurons gradually clear IgG, and our *in vitro* data illustrates that lysosomal degradation participates in this process. These results were validated *in vivo*, where inhibition of lysosomal proteases increased IgG retention in the CNS. Blocking lysosomal activity with E64d after SCI was well tolerated and produced a modest, although nonsignificant, trend toward improved locomotor recovery. However, inhibition of lysosomal activity did not completely prevent IgG clearance from the spinal cord. In this regard, we found that IgG injected into the CSF can rapidly reach the peripheral circulation via an Fc receptor-independent mechanism. Nevertheless, FcRn-KO mice exhibited improved locomotor recovery compared with WT mice after SCI. Collectively, these findings indicate that reducing IgG degradation prolongs antibody retention within the CNS without impairing functional recovery and may even promote recovery after SCI. These results suggest that targeting lysosomal and/or FcRn-mediated pathways could extend the persistence of infiltrating, non-autoimmune, and potentially beneficial natural antibodies in the injured CNS, while also enhancing the therapeutic efficacy of monoclonal antibody-based interventions following CNS injury.

The presence of IgG and IgM in the CNS during the intermediate and chronic phases of SCI is well documented and correlates with the emergence of antibody-secreting cells (ASCs) ^8,10^. Elevated circulating CNS-reactive antibodies have been consistently reported in murine models ^8,9^, as well as in the serum of patients with chronic SCI ^12,31-33^. However, these later-stage humoral mechanisms likely do not account for the antibodies detected in the injured spinal cord at the early time points examined here. At 1 hour and 1 day post-SCI, *de novo* antibody production is improbable, and Ankeny *et al.* observed no changes in serum IgG or IgM at 1 day after SCI, with alterations only emerging at later stages ^9^. Thus, the early immunoglobulin signal we observe most likely reflects acute barrier disruption and passive entry of circulating natural antibodies rather than a newly initiated humoral response, a conclusion further supported by our data showing that these antibodies are non-autoimmune.

To our knowledge, this study provides one of the first characterizations of immunoglobulin distribution during acute and subacute phases of SCI. We show that the three most abundant circulating antibodies, IgG, IgA and IgM, not only accumulate at and beyond the lesion site, but are also internalized by CNS-resident cells. Among immunoglobulins, IgG exhibited the widest distribution, likely reflecting its smaller molecular size (∼150 kDa vs. 900 kDa for IgM) and higher circulating levels in mice ^34^. Notably, IgG was detected in cervical segments as early as 1-hour post-SCI, with prominent accumulation around the central canal, suggesting partial dissemination through the CSF. Although this pattern implies a rostral movement of CSF after injury, no published studies have demonstrated a predominant rostral CSF flow within the central canal following SCI. Together, our observations on IgG movement along the central canal underscore the need to better understand CSF dynamics after SCI and to determine whether rostral dissemination of blood-derived molecules contributes to supraspinal pathology.

Microglia are known to express FcR and respond to IgG in diverse pathological contexts ^35^, and IgM can induce phagocytosis through complement receptor 3 ^36^, suggesting that microglia should readily engage with immunoglobulins. Surprisingly, we observed minimal microglia-immunoglobulin interaction during the acute and subacute phases of SCI. One explanation is that immunoglobulins enter the spinal cord via passive leakage rather than a structured humoral response early post-injury, limiting microglial engagement. In contrast, astrocytes robustly internalized immunoglobulins, consistent with earlier observations by Bernstein *et al*. showing IgG, IgM, and IgA colocalization with GFAP after SCI and cortical injury ^37,38^. Similarly, Fehlings and colleagues reported that exogenously administered human IgG to rats after cervical SCI interacted with astrocytes, but not microglia ^39^, although species-specific differences in IgG signaling should be considered. Given the diverse and context-dependent roles of astrocytes in CNS physiology and disease ^40^, understanding how immunoglobulin uptake influences their functions and shapes key astrocytic responses after SCI remains an important area for future research.

We found that IgG, IgM, and IgA rapidly infiltrate the spinal cord parenchyma after SCI and are predominantly internalized by neurons, indicating a strong neuron-immunoglobulin affinity. Neuronal uptake of immunoglobulins has been observed in multiple contexts, including IgG internalization by Purkinje cells *in vitro* ^5^, accumulation of autoantibodies in ventral horn neurons post-SCI ^8^, and neuronal IgG uptake following traumatic brain injury ^41^. Additional work has shown that IgG can modulate pain via FcR on dorsal root ganglion neurons ^7,42,43^, supporting neuron-immunoglobulin interactions as a conserved phenomenon across the nervous system. Notably, we observed immunoglobulin uptake by morphologically healthy neurons both at the lesion border and in distant segments, suggesting that antibodies may influence neuronal function far beyond the injury site. Our results show that, despite a large proportion of neurons internalizing immunoglobulins near the lesion border, these neurons do not appear to decrease in number within 1 week after SCI, suggesting that this interaction may not be neurotoxic. Supporting this results, IgG-based therapies have been explored in stroke, TBI, and SCI ^8,44-48^. After SCI, intravenous immunoglobulin (IVIg) from healthy donors reduces neuroinflammation, limits lesion progression, and improves functional recovery ^39,45,46^, and colocalizes with neuronal markers ^45^. *In vitro*, human IgG protects primary cortical neurons from ischemia-induced apoptosis ^44,49,50^, although the underlying mechanisms remain unclear. Importantly, these studies rely on human IgG applied to murine systems, raising the possibility that the reported benefits arise from interspecies interactions rather than physiological effects of endogenous IgG.

Despite the atrophic morphology of neurons in the ischemic zone, which likely reflects a stressed phenotype, IgG and IgM internalization was observed in neurons after MCAO. In contrast to SCI, statistically significant neuronal uptake of IgA was not observed in this model. Following MCAO, neurons in the lesion core undergo rapid necrotic death, whereas neuronal loss in the surrounding penumbra progresses more gradually ^51,52^. Although many penumbral neurons display features of neurodegeneration, some remain sufficiently viable to activate pro-survival pathways. For example, Shibata *et al*. reported early Akt phosphorylation in penumbral neurons during the early hours after transient MCAO ^53^, consistent with established roles for Akt signaling in neuronal survival ^54^. The presence of phosphorylated Akt suggests a potential therapeutic window for neuronal rescue after stroke. While the functional consequences of IgG or IgM uptake remain uncertain, these findings highlight possible therapeutic avenues, particularly the design of monoclonal antibody-based strategies capable of neuronal internalization to modulate intracellular death pathways and enhance neuroprotection.

Intra-cisterna magna administration of Alexa488-conjugated IgG from healthy mouse serum into the CSF of recipient mice led to rapid uptake by neurons and astrocytes, indicating that immunoglobulin internalization is not solely an injury-driven phenomenon. This observation is consistent with a recent report of neuronal IgG uptake following intrastriatal injection in healthy mice ^55^. However, because tissue-directed injections such as intrastriatal delivery may introduce local microdamage through needle penetration and trigger injury-related signaling ^56^, the use of an i.c.m. approach here provides clearer evidence that non-injured neurons and astrocytes can internalize IgG under physiological conditions.

Several mechanisms have been proposed to explain how IgG enters cells. For example, antigen-presenting cells internalize IgG through FcR-mediated uptake of immune complexes ^57,58^. Similarly, the therapeutic anti-PD-L1 IgG monoclonal antibody avelumab as well as the anti-CD20 IgG monoclonal antibody rituximab, both used in cancer treatment, are cleared by circulating immune cells via FcγR-mediated internalization ^59,60^. Ward *et al.* suggested that endothelial cells may internalize IgG non-selectively through fluid-phase pinocytosis ^24^. In a study using an *in vitro* BBB model derived from human induced pluripotent stem cells, the addition of amiloride, a micropinocytosis blocker, was found to decrease IgG uptake by nearly half ^61^. Despite evidence that FcR-dependent mechanisms can mediate IgG trafficking, our data indicate that Fc fragments play little role in neuronal IgG uptake. Indeed, neurons internalize F(ab’)₂ fragments as efficiently as full-length IgG following i.c.m. administration, suggesting that uptake is largely independent of Fc-FcR interactions. Moreover, because Fc fragments are smaller than F(ab’)₂ fragments, their poor uptake argues against passive mechanisms such as pinocytosis and instead supports an Fc-independent, selective endocytic process.

Confocal colocalization studies and the progressive decline in neuronal IgG levels over time demonstrate that neurons can internalize and clear immunoglobulins. Using an immortalized human endothelial cell line, Ward *et al*. reported that intracellular IgG not recycled to the cell surface is routed to lysosomal degradation ^24^. Consistent with this, we show both *in vitro* and *in vivo* that neurons also clear internalized IgG, and that this process is partially attenuated by E64d, an irreversible, membrane-permeable cysteine protease inhibitor that primarily targets cathepsins ^62^, key enzymes involved in lysosomal protein degradation. Notably, although E64d treatment increases neuronal IgG accumulation, clearance still occurs. Several explanations are possible. First, the inhibitor dose may have been insufficient to reach the CNS is quantities needed to fully block lysosomal activity. Second, because the inhibition is irreversible, newly formed lysosomes produced after E64d exposure and washout may retain functional protease activity. Third, IgG may also undergo degradation through lysosome-independent pathways, a possibility particularly relevant in neurons, where the ubiquitin-proteasome system constitutes a major degradation route distinct from lysosomal proteolysis ^63^.

Monoclonal antibody-based therapies targeting CNS antigens face several obstacles, with efficient clearance being a major challenge (see review ^64^). Because systemic delivery is hindered by blood-CNS barriers, intrathecal administration is often favored to deliver antibodies directly into the CSF ^65^. However, this strategy remains suboptimal, as antibody clearance begins within minutes and results in a short CSF half-life ^66-68^. This issue is broadly relevant across CNS disorders and may be particularly limiting in SCI. One of the most promising therapies in this context, blocking the neuron-enriched Nogo-A protein using anti-Nogo antibodies, has shown robust preclinical efficacy in promoting axonal regeneration and functional recovery ^69,70^. Yet, a recently published phase 2b clinical trial in acute cervical SCI found no significant improvement in the primary endpoint (upper-extremity motor score) between intrathecally treated patients and placebo ^71^. Notably, the antibody displayed a short ∼10-hour half-life, and among motor-incomplete patients, higher CSF antibody levels at Day 5 correlated with better outcomes. Rapid clearance after intrathecal delivery has also been documented in patients with CNS lymphoma treated intraventricularly with Rituximab ^72^. Interestingly, our findings suggest that E64d pretreatment may prolong the intracellular presence of exogenously administered IgG in CNS-resident cells, including neurons. Whether this expanded intracellular IgG pool retains antigen recognition specificity and functional activity remains unknown. If similar effects occur with monoclonal antibodies directed against intracellular or CNS-restricted antigens, E64d could potentially extend their therapeutic window, a possibility that warrants further investigation.

Another advantage of E64d is its ability to cross CNS barriers. Furthermore, this compound has completed a phase III clinical trial for Duchenne muscular dystrophy, where it was found to be safe and well tolerated in humans, although no therapeutic benefit was observed^73^. E64d has also been evaluated in several animal models of CNS disorders, including Alzheimer’s disease ^74,75^, stroke ^76,77^, and SCI ^78,79^, where it has been reported to exert neuroprotective effects. In our study, acute E64d administration produced a modest, though not statistically significant, improvement in locomotor recovery. Although the acute benefit was small and delayed treatment showed no effect, our primary aim was to assess whether E64d could serve as an adjuvant to monoclonal antibody-based therapies. Taken together, its lack of detrimental effects, ability to cross CNS barriers, and established safety profile in humans positions E64d as a promising candidate to investigate whether it can enhance the bioavailability and efficacy of monoclonal antibody therapies for CNS injury and other neurological disorders.

While the blockade of the lysosomal pathway did not fully prevent IgG clearance *in vitro* and produced only a modest enhancement of IgG retention in the CNS *in vivo*, these observations suggest that additional mechanisms likely contribute to IgG degradation or recycling. One such pathway involves FcRn, which binds monomeric IgG in the acidic environment of endosomes and mediates its recycling back to the cell surface ^80^. Beyond its canonical recycling role, FcRn is abundantly expressed in the syncytiotrophoblast layer of the placenta, where it facilitates transcytosis of maternal IgG to the fetus and enables passive immunity ^28,81-83^. Cooper *et al*. showed that antibodies with reduced affinity for FcRn persist longer in the CNS and display lower serum levels in comparison with those with higher affinity for the receptor following intra-cranial administration in rats^27^. Given that FcRn expression has also been identified in brain endothelial cells ^29,84^, this raises the possibility that FcRn participates in IgG efflux from the CNS. However, our results diverge from previous reports, showing that mice lacking FcRn transfer (rabbit) IgG to the circulation in quantities comparable to those observed in WT mice. These findings underscore the importance of elucidating the mechanisms governing IgG externalization from the CNS, as this process could be strategically leveraged either to enhance the efficacy of antibody-based treatments or to reduce CNS levels of self-reactive antibodies in autoimmune diseases such as systemic lupus erythematosus, as described in a companion study (Laroche *et al*., unpublished data). A recent study suggests that these mechanisms may involve the Integral Membrane Protein 2A (ITM2A), a membrane protein enriched in brain endothelial cells that has been proposed to mediate receptor-dependent transcytosis ^85^. However, current evidence supporting its role is limited to a single *in vitro* study, leaving its contribution to IgG transport across the BBB *in vivo* uncertain.

To our knowledge, this is the first examination of FcRn-KO mice in a CNS injury models such as SCI, where we demonstrate a significant improvement in locomotor recovery. However, FcRn-KO mice present notable baseline physiological differences compared with WT mice, as FcRn mediates the recycling of both IgG and albumin ^86,87^. Accordingly, FcRn-KO mice display reduced circulating levels of both proteins under normal conditions ^88,89^. Because contusive SCI causes vascular rupture and the infiltration of blood-derived components into the parenchyma, the reduced albumin load in FcRn-KO mice could conceivably contribute to a reduction in albumin extravasation. The effects of albumin extravasation on SCI pathophysiology remain poorly defined, making it difficult to determine whether the enhanced functional recovery observed here stems from the absence of FcRn itself or from secondary consequences of this genetic modification. Despite these uncertainties, our findings highlight the therapeutic potential of modulating immunoglobulins clearance pathways in CNS injury. They identify the FcRn axis as a promising target and support future studies aimed at dissecting its mechanistic contribution to neuroprotection and recovery. This beneficial effect of FcRn inhibition is consistent with findings demonstrating that FcRn antagonists have therapeutic potential in neuroautoimmune diseases, with FcRn-targeting therapies already approved clinically for myasthenia gravis ^90^.

In conclusion, this study identifies neurons as major cellular targets for immunoglobulins within the CNS under both physiological conditions and after injury. We demonstrate that neurons constitutively internalize IgG through an Fc-independent mechanism and that this process is strongly enhanced following CNS injury, when blood-derived immunoglobulins gain access to the parenchyma. Once internalized, IgG is actively cleared by neurons, with lysosomal degradation representing a principal elimination pathway. In addition, our data show that loss of FcRn leads to increased IgG retention within the CNS without negatively affecting functional recovery after SCI. Together, these findings establish neuronal uptake, lysosomal degradation, and FcRn-dependent handling as key determinants of immunoglobulin dynamics in the CNS, and provide a cellular framework for understanding how antibodies are processed and eliminated within neural tissue.

## MATERIALS AND METHODS

### Mice

Both male and female mice were included in this study. Adult (2-4 months old) C57BL/6 and B6;129S mice were obtained from Charles River Laboratories or The Jackson Laboratory (JAX). FcγR-KO mice were purchased from JAX (stock #002847) and subsequently backcrossed for at least 7 generations onto the C57BL/6J background at the Animal Research Facility of the Centre de recherche du Centre hospitalier universitaire (CRCHU) de Québec– Université Laval. Accordingly, C57BL/6J mice were used as control animals for this strain. FcγRIIb-KO mice on the B6;129S background were obtained from JAX (stock #002848), and B6.129S were used as controls. *Cx3cr1*^CreER^ mice were sourced from the European Mouse Mutant Archive, following prior authorization from Dr. Steffen Jung (Rehovot, Israel), and genotyped according to previously established protocols ^91^. Breeders for *Emx1*^cre^ knock-in mice (stock #005628, ^92^), *Rosa26-tdTomato* (*R26*^tdT^) reporter mice (also known as Ai9; stock #007905, ^93^), and FcRn-KO mice (stock #003982, ^89^), all on the C57BL/6J background, were purchased from JAX. *Cx3cr1*^CreER^, *Emx1*^cre^, Ai9, and FcRn-KO lines were maintained and bred locally at the Animal Facility of the CRCHU de Québec–Université Laval. All mice were housed in individually ventilated cages supplied with HEPA-filtered air (30-70 air exchanges per hour) under a 12-hour light/dark cycle, with unrestricted access to food and water. Environmental conditions were maintained at 23 ± 2 °C with a relative humidity of 50 ± 5%. All experimental procedures were approved by the *Comité de protection des animaux de l’Université Laval* (protocols #CHU-22-1136, #CHU-23-1239, and #CHU-24-1635) and conducted in compliance with relevant ethical regulations and guidelines of the Canadian Council on Animal Care.

### Tamoxifen treatment

To induce cre-mediated recombination in *Cx3cr1*^CreER^::*R26*^tdT^ mice, animals were orally administered 10 mg of tamoxifen dissolved in a 1:10 ethanol/corn oil solution. Mice received two gavages 48 hours apart at postnatal day (P) 30 and P32.

### Spinal cord injury

Mice were anesthetized with isoflurane and subjected to a thoracic laminectomy performed at vertebral levels T9-T10, corresponding to spinal cord segments T10-T11. Following stabilization of the vertebral column, a moderate contusive SCI was induced using the Infinite Horizon impactor (Precision Systems & Instrumentation) set to deliver a force of 50 kdyn. After injury, the paravertebral musculature was sutured, and the skin was closed using surgical staples. During the postoperative period, animals received manual bladder expression twice daily to ensure urinary voiding and reduce the risk of urinary complications.

### Middle cerebral artery occlusion

Transient focal cerebral ischemia was induced in adult male mice (3-4 months old) under isoflurane anesthesia using the intraluminal filament MCAO model, following previously established protocols ^94^. Briefly, mice received a subcutaneous injection of sustained-release buprenorphine (1 mg/kg) for pre-emptive analgesia 1 hour prior to surgery. The cervical region was shaved and locally anesthetized with a subcutaneous injection of lidocaine/bupivacaine (3 mg/kg). A midline cervical incision was made, and the left common carotid artery was exposed and ligated under a surgical microscope. A temporary microvascular clip was applied to the internal carotid artery, after which a silicone-coated 7-0 nylon monofilament (Doccol Corporation) was inserted into the internal carotid artery and advanced until it reached the origin of the middle cerebral artery to induce ischemia. Mice underwent a 45-minute occlusion of the middle cerebral artery. Throughout the procedure, body temperature was continuously monitored and maintained between 36-37 °C using a feedback-controlled heating system (Harvard Apparatus). For perioperative support, mice were administered carprofen (10 mg/kg, s.c.) for analgesia and 1 mL of Ringer’s lactate subcutaneously to prevent dehydration. At the end of surgery, mice received an additional 0.5 mL of Ringer’s lactate and were transferred to a heated recovery cage. During the first 5 postoperative days, mice received supportive hydration with Ringer’s lactate (1 mL, s.c.) twice daily. Postoperative analgesia was maintained with daily carprofen injections (10 mg/kg, s.c.) for the first 3 days, complemented by sustained-release buprenorphine (1 mg/kg, s.c.) on postoperative days 2 and 4.

### Intra-cisterna magna injections

Mice were injected i.c.m. with either Alexa Fluor^®^ 488 ChromPure^®^ Mouse IgG whole molecule (20µg/µL diluted in PBS, 5 µL injected per mouse; Jackson ImmunoResearch, catalog #015-540-003), Alexa Fluor^®^ 488 ChromPure^®^ Mouse IgG F(ab’)₂ fragment (20µg/µL diluted in PBS, 5 µL injected per mouse; Jackson ImmunoResearch, #015-540-006), Fluorescein (FITC) ChromPure® Mouse IgG Fc fragment (20µg/µL diluted in PBS, 5 µL injected per mouse; Jackson ImmunoResearch, #015-090-008), ChromPure® Rabbit IgG whole molecule (40µg/µL diluted in PBS, 5 µL injected per mouse; Jackson ImmunoResearch, catalog #011-000-003) or PBS alone (5 µL/mouse). The i.c.m. treatment consisted of a single injection using a pulled-glass micropipette connected to a 10-µL Hamilton syringe.

### Tissue processing

Mice were euthanized by overdose with ketamine (400 mg/kg) and xylazine (40 mg/kg), followed by transcardial perfusion with phosphate-buffered saline (PBS) and 1% paraformaldehyde (PFA; pH 7.4) in PBS. Spinal cords were carefully dissected from the vertebral column, post-fixed for an additional 48 hours in 1% PFA at 4°C, and subsequently cryoprotected by immersion in PBS containing 20% sucrose for at least 24 hours prior to sectioning. For experiments involving i.c.m. injection of immunoglobulins and specific IgG fragments (i.e., Fc and F(ab’)_2_), spinal cords were cut into 4-mm blocks corresponding to the upper cervical, mid-thoracic, and upper lumbar regions. In SCI experiments, a 12-mm segment centered on the lesion site was collected and subdivided into three equal portions corresponding to the lesion epicenter, as well as rostral and caudal regions, as previously described ^95^. For each animal, the three spinal cord segments were embedded in Shandon™ M-1 Embedding Matrix (Thermo Fisher Scientific). Tissue sections were cut at 14 µm using a cryostat (CM3050S; Leica Biosystems) and mounted directly onto permanently positively charged Surgipath X-tra^®^ glass slides (Leica Microsystems). Sections were stored at -20 °C until further processing.

Brain tissue was sectioned coronally at 30 µm using a sliding-blade microtome (SM2010; Leica Biosystems). Free-floating sections were transferred to an antifreeze storage solution consisting of 12.5% sodium phosphate buffer (prepared from 0.05 M sodium phosphate monobasic and 0.154 M sodium phosphate dibasic in Milli-Q water), 37.5% diethyl pyrocarbonate-treated water, 30% ethylene glycol, and 20% glycerol (all reagents from Thermo Fisher Scientific). Brain sections were stored in this solution at -20 °C until further processing.

### Immunostaining

Immunofluorescence staining was performed as described previously ^95^. Primary antibodies used in this study were obtained from the sources listed below (catalog numbers indicated) and applied at the specified dilutions for slide-mounted tissue sections. For free-floating brain sections, both primary and secondary antibodies were used at twice the concentration applied to slide-mounted sections: rat anti-CD31 (1:1,000, BD Biosciences, catalog #557355), rabbit anti-Iba1 (1:500, FUJIFILM Wako, 019-19741), goat anti-IgA (1:250, Thermo Fisher Scientific, 62-6700), Alexa488-conjugated goat anti-IgG (1:125, Jackson ImmunoResearch, 115-545-164), goat anti-IgM (1:250, Jackson ImmunoResearch, 115-005-020), rabbit anti-laminin (1:1,500, Dako, Z0097), rabbit anti-NeuN (1:1,000, Cell Signaling Technology, 12943), rabbit anti-RFP/tdT (1:500, Rockland Immunochemicals, 600-401-379), and guinea pig anti-S100β (1:500, Synaptic Systems, 287004). Alexa Fluor secondary antibody conjugates (1:500, Thermo Fisher Scientific) were used as secondary antibodies. Nuclear counterstaining was performed using 4’,6-diamidino-2-phenylindole, dilactate (DAPI; 1 μg/ml, Thermo Fisher Scientific).

### Image acquisition and quantification

For quantification of the area occupied by IgG, IgA, and IgM immunostaining in the injured spinal cord, images were acquired at 20× magnification using a Zeiss AxioScan.Z1 slide scanner equipped with a 20× Plan-Apochromat objective (NA 0.8) and ZEN 2.3 (Blue edition) software. Signal-positive area was quantified in MATLAB R2020a (MathWorks) by calculating the number of pixels exceeding a predefined threshold and expressed as a percentage of the total tissue area, which corresponded to the entire coronal section. Quantification of tdT^+^ microglia or S100β^+^ astrocytes colocalizing with IgG, IgM, or IgA in the spinal cord white matter was performed on 20× AxioScan.Z1 images using ImageJ with the Cell Counter plugin (NIH). NeuN^+^ neurons colocalizing with immunoglobulins were quantified directly at the microscope in the spinal cord gray matter using a Nikon Eclipse 80i equipped with a Color QImaging QIClick camera (01-QIClick-R-F-CLR-12) at 40× magnification, and BIOQUANT Life Science software (v. 17.5, Bioquant Image Analysis Corporation). In coronal brain sections from MCAO mice, the lesion area was defined as the region exhibiting reduced NeuN signal intensity compared with the contralateral hemisphere. Within this lesion area, NeuN^+^ neurons colocalizing with immunoglobulins were counted. Only immunolabeled tdT^+^, S100β^+^, or NeuN^+^ cells containing a DAPI-stained nucleus were counted, and results were expressed as cells per mm².

For cell culture experiments, 10× images were acquired using a Zeiss AxioScan.Z1. Quantitative analysis was performed in MATLAB R2020a by thresholding the RFP channel to generate a mask defining RFP^+^ regions, within which mean IgG fluorescence intensity was measured and normalized to values obtained from negative control samples.

For analysis of Alexa488-conjugated IgG administered i.c.m., neurons colocalizing with IgG were counted in the CoA region and spinal cord gray matter using BIOQUANT Life Science software, as described above. The area occupied by the IgG signal in the spinal cord white matter was quantified in MATLAB using 10× AxioScan.Z1 images. For E64d-treated mice, two independent experiments were analyzed separately using experiment-specific thresholds, and data were normalized to the mean IgG-covered area of vehicle-treated controls within each experiment.

Representative images were selected from animals with values at or near the group mean. Full brain sections were imaged at 2.5× magnification, full spinal cord sections at 10×, and regions showing cellular labeling at 20× using a Zeiss LSM 800 confocal microscope (405, 488, 561, and 640 nm lasers) equipped with a Zeiss Axiocam 506 Mono camera. Mosaics were assembled in ZEN 2.3, and Z-stacks were processed in ImageJ (NIH) to generate maximum-intensity projections. The neuron cell culture video was acquired on the same confocal microscope at 40× magnification, and three-dimensional reconstruction was performed using Imaris software (Bitplane, Oxford Instruments).

### Three-dimensional reconstruction

A representative confocal z-stack was acquired using a Zeiss LSM 800 confocal microscope (405, 488, 561, and 640 nm lasers) equipped with a Zeiss Axiocam 506 Mono camera. Images were collected with a 40× objective at 0.2 µm z-intervals. The stack was imported into Imaris software (Bitplane, Oxford Instruments) to generate a three-dimensional rendering. The final output corresponds to a representative 3D reconstruction video generated from the acquired z-stack.

### Bio-dot analysis

#### Blood collection and plasma preparation

Following SCI or sham surgery, whole blood was collected from anesthetized mice via terminal cardiac puncture using a 22-gauge syringe and transferred into EDTA-coated tubes (Sarstedt) to prevent coagulation. Samples were kept on ice and processed immediately to obtain plasma. Blood was centrifuged to separate the antibody-containing plasma fraction, which was then collected, aliquoted, and stored at -80°C until analysis. Serum samples from lupus mice, provided by Dr. Eric Boilard, were included as positive controls.

#### Spinal cord and brain lysate preparation

Following blood collection, mice were transcardially perfused with ice-cold PBS to remove intravascular blood. Spinal cords from naïve and SCI mice, as well as brains from naïve mice, were rapidly dissected. Tissues were homogenized in a non-denaturing lysis buffer containing: 50 mM Tris-HCl, 150 mM NaCl, 2 mM EDTA, 10% Glycerol, 1 mM phenylmethanesulfonyl fluoride (PMSF), protease inhibitor cocktail (Sigma-Aldrich, catalog #P8340), and two phosphatase inhibitor cocktails (Sigma-Aldrich, catalog #P5726 and #P0044). Lysates were clarified by centrifugation, and supernatants were collected and stored at -80 °C until analysis. In subsequent steps, spinal cord lysates were used as a source of primary antibodies to determine whether infiltrated immunoglobulins exhibited self-reactivity, while brain lysates served as the source of CNS antigens for spotting onto the membranes.

#### Bio-dot immunoassay

Brain lysates were applied at 5 µg, 2 µg or 1 µg onto 0.2-µm nitrocellulose membranes (Amersham Cytiva) using a Bio-Dot Microfiltration Apparatus (Bio-Rad), allowing antigen binding by gravity filtration. Membranes were then blocked with 1% bovine serum albumin (BSA; Sigma-Aldrich) prepared in TBS, followed by incubation with plasma, serum or spinal cord lysate samples diluted 1:200 for plasma and serum or 1:100 for spinal cord lysates in TBS containing 1% BSA and 0.05% Tween-20 (Sigma-Aldrich). After sample incubation, membranes were probed with goat horseradish peroxidase (HRP)-conjugated anti-mouse secondary antibody (1:2,000 dilution, Jackson ImmunoResearch, catalog #115-035-003). Immunoreactive signals were developed using Immobilon^®^ Forte Western HRP Substrate (Merck Millipore). Negative controls were included by omitting brain lysates during antigen application. Uncropped and unprocessed scans of immunoblots are shown in Supplementary Figure 2.

#### Quantification

Signal intensity was quantified in ImageJ by measuring the mean gray value of wells corresponding to the 5-µg antigen load for each sample, after subtracting the corresponding negative control signal.

### Primary neuron culture

Primary cortical neurons were prepared from P0 *Emx1*^Cre^::*R26*^tdT^ mouse pups. Cerebral cortices were dissected in ice-cold Hibernate A-based medium (Gibco) supplemented with 0.5× B27 (Gibco), 0.5× L-glutamine (Gibco), and penicillin/streptomycin (Life Technologies). Meninges and blood vessels were carefully removed, and the tissue was enzymatically dissociated using 0.25% Trypsin (Gibco) in the presence of DNase I. The tissue was then gently triturated to obtain a single-cell suspension, filtered through a 70-µm cell strainer, and centrifuged. Cells were resuspended in plating medium consisting of Neurobasal A (Gibco) supplemented with 10% heat-inactivated FBS (Wisent), 1× B27, 1× N-2 (Gibco), 1× penicillin/streptomycin, 1x HEPES (Gibco), and 1x GlutaMAX (Gibco). Two hours after plating, the medium was replaced with growth medium composed of Neurobasal A supplemented with 1× B27, 1× N-2, 1× penicillin/streptomycin, 0.25× L-glutamine, and 0.25× GlutaMAX. Neurons were seeded onto poly-D-lysine-coated glass coverslips in 24-well plates and maintained at 37°C in a humidified incubator with 5% CO. Half of the culture medium was replaced every three days. Neurons were used for experiments after 7 days *in vitro*.

### IgG incubation

Primary neurons were incubated with ChromPure^®^ Mouse IgG whole molecule (10 µg/ml; Jackson ImmunoResearch, catalog # 015-000-003) diluted in growth medium for 48 hours. At the end of the incubation period, neurons were fixed with 4% PFA for 1 hour at 37°C and processed for immunostaining and analysis. To assess IgG degradation over time, parallel experiments were performed in which the IgG-containing medium was removed after the initial 48-hour incubation period and replaced with fresh IgG-free medium. Neurons were then fixed 1, 4, or 24 hours after medium replacement and processed for immunostaining and analysis.

### Lysosomal protease inhibition using E64d

#### In vitro

To pharmacologically inhibit lysosomal activity in neurons exposed to IgG for 48 hours, the lysosomal cysteine protease inhibitor E64d (40 µg/ml in ethanol, Tocris Bioscience, catalog #4545) or vehicle (ethanol at the same final concentration) was added to the growth medium during the final 24 hours of the 48-hour IgG incubation period. At the end of the 48-hour incubation, the medium was removed and replaced with fresh IgG- and E64d-free medium. Neurons were maintained under inhibitor-free conditions for 4 or 24 hours before fixation with 4% PFA and subsequent immunostaining and analysis.

#### In vivo

Mice received an i.p. injection of E64d (7.5 mg/kg), prepared in PBS containing 1.5% dimethyl sulfoxide (DMSO), one hour prior to i.c.m. administration of IgG or vehicle prepared in PBS containing 1.5% DMSO. Animals were subsequently processed at 1 and 4 hours after i.c.m injection for tissue collection. E64d was also administered i.p. following SCI. mice received E64d either acutely (1 dpi) or in a delayed manner (14 dpi), according to the experimental design. Animals were evaluated using BMS behavioral analysis as described below.

### ELISA

Rabbit IgG levels in mouse plasma were quantified using the Rabbit IgG ELISA Kit (Abcam, catalog #ab187400), following the manufacturer’s instructions. Briefly, plasma samples were collected at 4 hours after i.c.m. injection of rabbit IgG, as explained in the *Blood collection and plasma preparation* subsection. Samples were diluted to 1:10,000 and added to microplate wells pre-coated with anti-rabbit IgG antibodies. After incubation and washing steps, an HRP-conjugated detection antibody was applied, followed by substrate development. Absorbance was measured at 450 nm using a microplate reader, and IgG concentrations were determined by interpolation from a standard curve generated with known concentrations of rabbit IgG.

### Behavioral analysis

Locomotor function after SCI was assessed in an open field using the BMS, as originally described by Basso *et al*. ^96^. Prior to behavioral assessment, all experimental groups were verified to exhibit comparable impact forces and equivalent spinal cord displacement at the time of injury. Behavioral scoring was performed by two observers blinded to genotype and experimental conditions.

### Statistical analysis

Statistical evaluations were performed using two-way ANOVA or repeated-measures ANOVA where appropriate. The multiple comparisons adjustment was made using the Bonferroni correction. All statistical analyses were performed using the GraphPad Prism software (v. 10.6.0).

## ACKNOWLEDGEMENTS

This study was supported by grants from the Canadian Institutes of Health Research (CIHR; (PJT-197909 to S.L. and PJT-124998 to E.B. and S.L.) and the Wings for Life Spinal Cord Research Foundation (WFL-CA-18/24, Project # 316 to S.L.). This work was also supported by the Fonds de recherche du Québec (FRQ) through a Research Centre grant to the CRCHU de Québec–Université Laval (Grant # 30641). A.B. and D.B. are supported by studentship awards from the FRQ–Santé (FRQS). A.L. is recipient of a studentship award from The Arthritis Society. N.B is a Senior research Scholar from the FRQS. E.B. is supported by a Merit salary award from the FRQS. During the preparation of this manuscript, the authors used Microsoft 365 Copilot to assist with grammar correction and readability improvements. All content was subsequently reviewed and edited by the authors, who take full responsibility for the final version of the published article.

## AUTHOR CONTRIBUTIONS

**Adrian Castellanos-Molina**: Conceptualization, Methodology, Formal Analysis, Investigation, Data Curation, Writing – Original Draft, Writing – Review & Editing, Visualization. **Ana Boisvert:** Conceptualization, Methodology, Formal Analysis, Investigation, Writing – Original Draft, Writing – Review & Editing. **Juliette Ferry**: Formal Analysis, Investigation. **Dominic Bélanger**: Formal Analysis, Investigation. **Audrée Laroche**: Conceptualization, Writing – Review & Editing. **Romain Menet**: Methodology, Investigation. **Frédérique Crépeau**: Formal Analysis, Investigation. **Martine Lessard**: Methodology, Investigation. **Nadia Fortin**: Formal Analysis, Investigation. **Nicolas Vallières**: Methodology, Formal Analysis, Investigation, Data Curation, Visualization. **Isabelle Allaeys:** Methodology, Investigation. **Nicolas Bertrand**: Resources, Writing – Review & Editing. **Ayman ElAli**: Methodology, Resources, Writing – Review & Editing. **Éric Boilard**: Conceptualization, Resources, Writing – Review & Editing. **Steve Lacroix**: Conceptualization, Formal Analysis, Resources, Data Curation, Writing – Original Draft, Writing – Review & Editing, Supervision, Project administration, Funding acquisition.

