## Supplemental Figures 1-8 for "Immunoglobulins are rapidly internalized by neurons after CNS injury and cleared through lysosomal degradation"

### SUPPLEMENTARY FIGURES

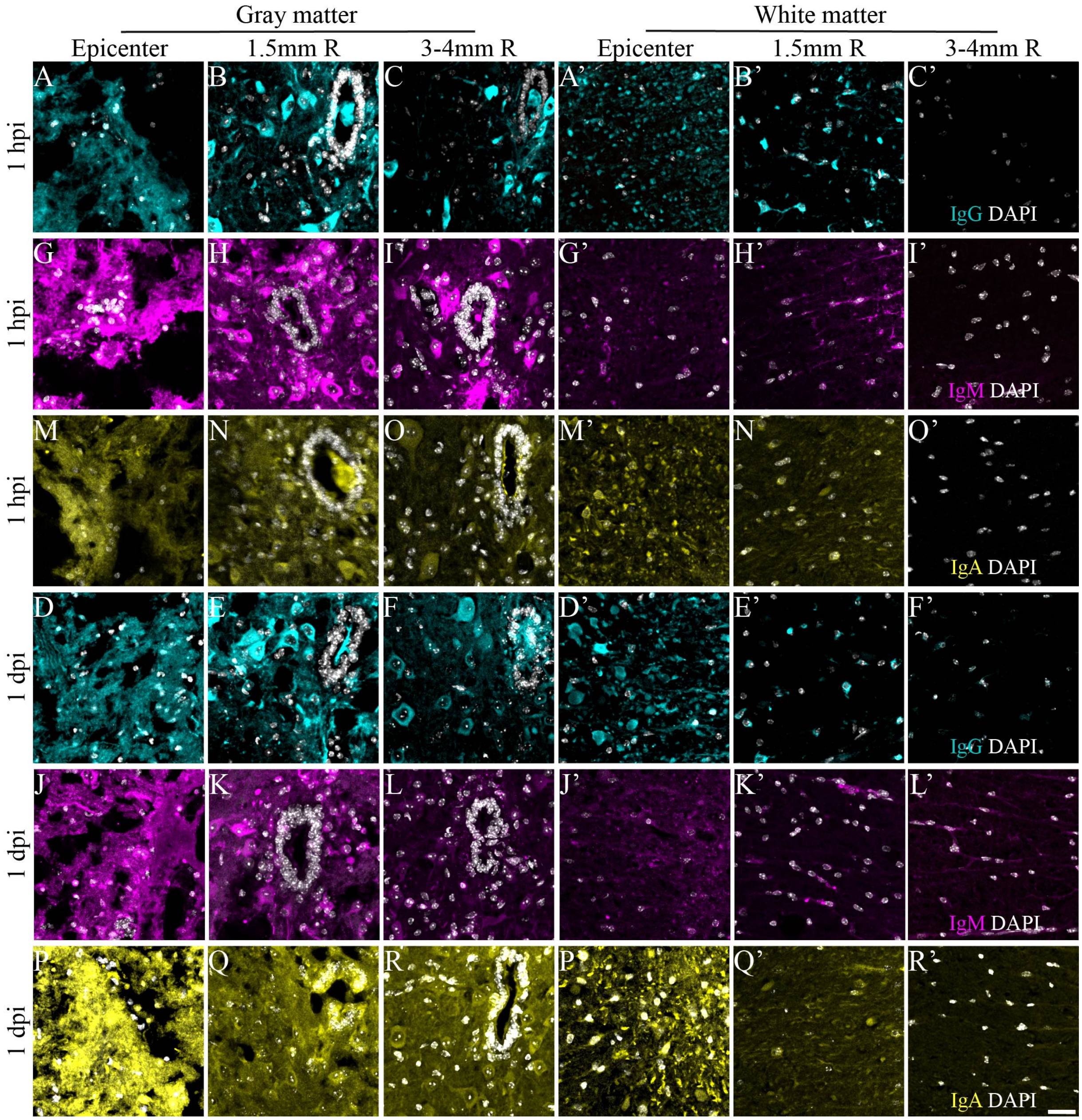


**Supplementary Figure 1. Spatial distribution and cellular localization of immunoglobulins in spinal cord gray and white matter after injury.** Representative confocal images of IgG, IgM, and IgA immunofluorescence staining in gray and white matter at different rostral levels and timepoints following SCI. **A**-**F**, IgG (cyan) and DAPI (grey) staining in gray matter at the lesion epicenter (**A**, **D**) 1.5 mm rostral (**B**, **E**), and 3-4 mm rostral (**C**, **F**) at 1 hour (**A**-**C**) and 1 day (**D**-**F**) post-SCI. **G**-**L**, IgM (magenta) and DAPI (grey) staining at the same anatomical levels at 1 hour (**G**-**I**) and 1 day (**J**-**L**). **M**-**R**, IgA (yellow) and DAPI (grey) staining at the same anatomical levels at 1 hour (**M**–**O**) and 1 day (**P**–**R**). **A′**-**R′**, Images of each immunoglobulin in white matter at the same anatomical regions and timepoints. Scale bars: 25 μm (shown in **F**, **F′**, **L**, **L′**, **R** and **R′**).

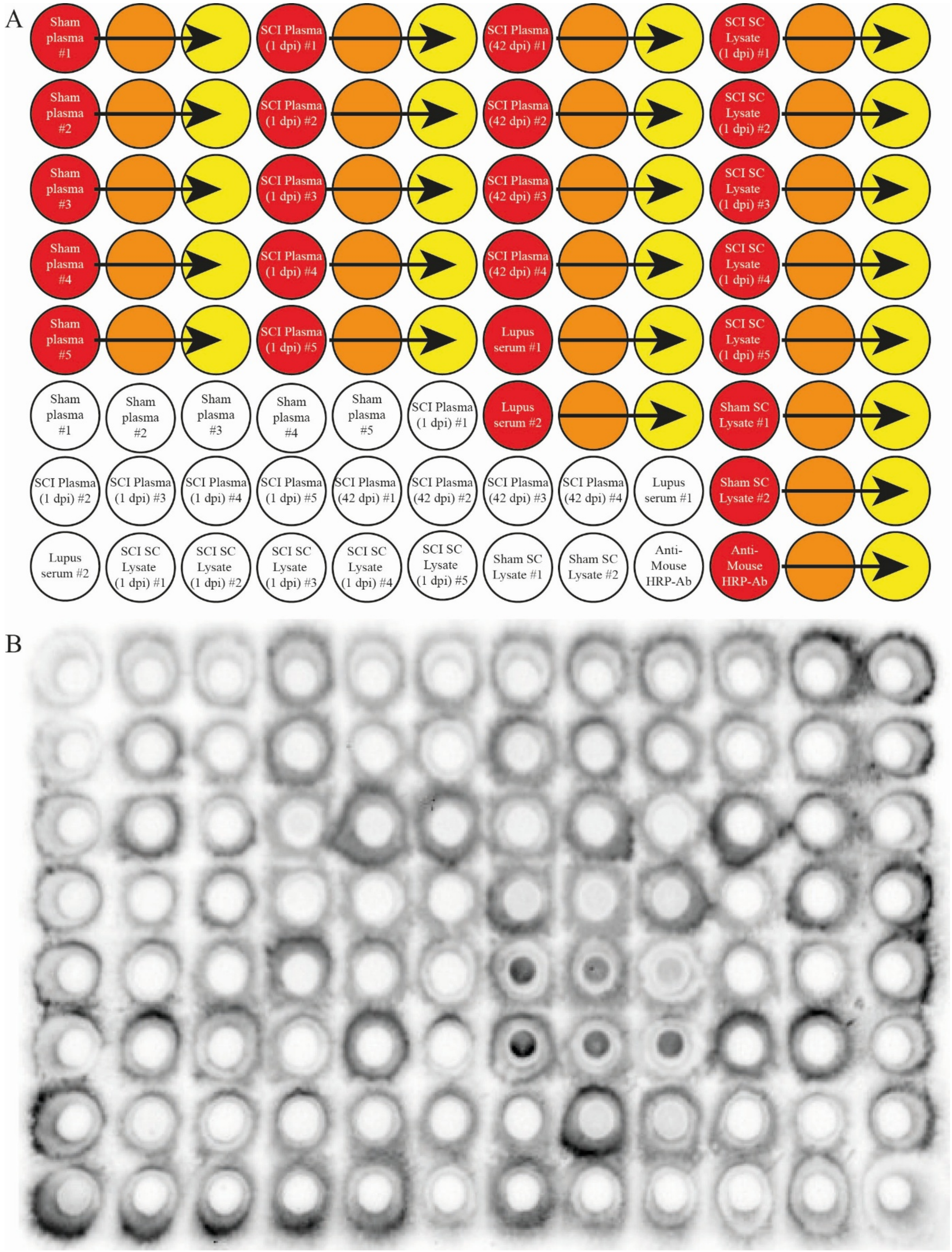


**Supplementary Figure 2. Immunoglobulins derived from serum and spinal cord lysate during the acute phase of SCI do not significantly recognize CNS antigens.** **A**, Schematic showing the samples loaded into each well of the bio-dot immunoassay. **B**, Image of the full membrane. Dark signal confined within a well indicates immunoreactivity of plasma- or spinal cord-derived antibodies to CNS antigens. Serum from lupus mice was included as a positive control, as their autoantibodies are known to recognize CNS antigens.

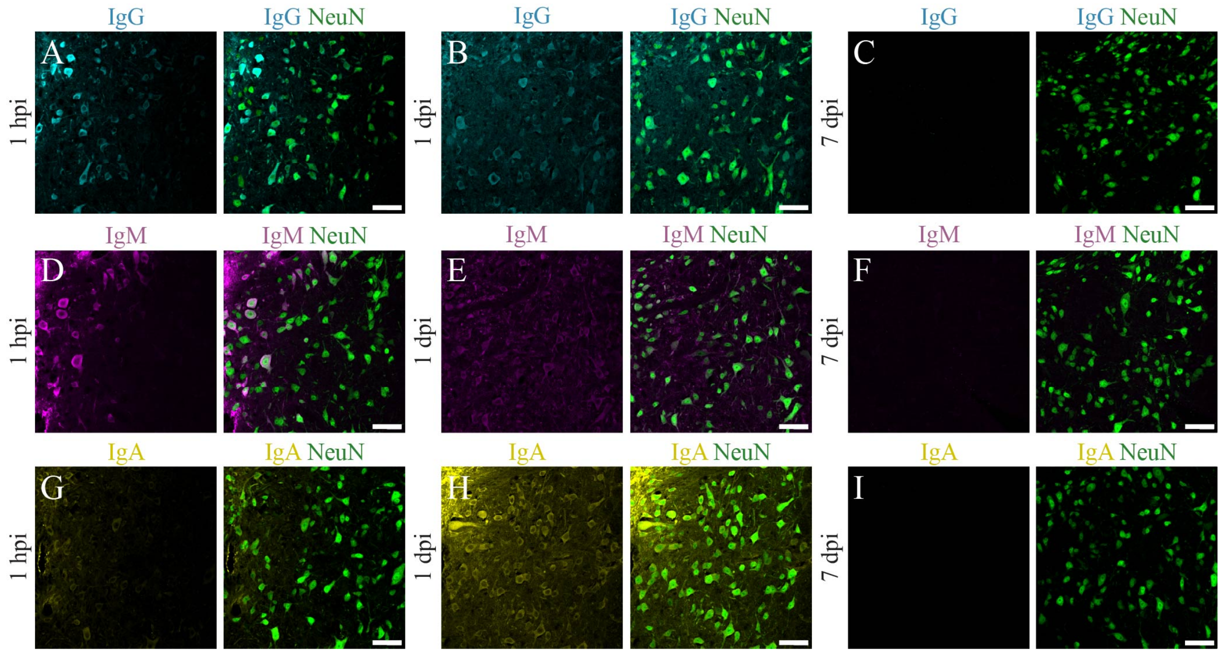


**Supplementary Figure 3. Immunoglobulins are internalized within neurons during the acute and subacute phases of SCI. A-C**, Representative confocal images showing IgG (cyan) within NeuN$^+$ neurons (green) at 1 hour (**A**), 1 day (**B**), and 7 days (**C**) post-SCI at 3-4 mm rostral to the lesion epicenter. **D-F**, IgM (magenta) within NeuN$^+$ neurons at the same location and timepoints. **G-I**, IgA (yellow) within NeuN$^+$ neurons at the same location and timepoints. Scale bars: 50 µm (shown in the merged panels).

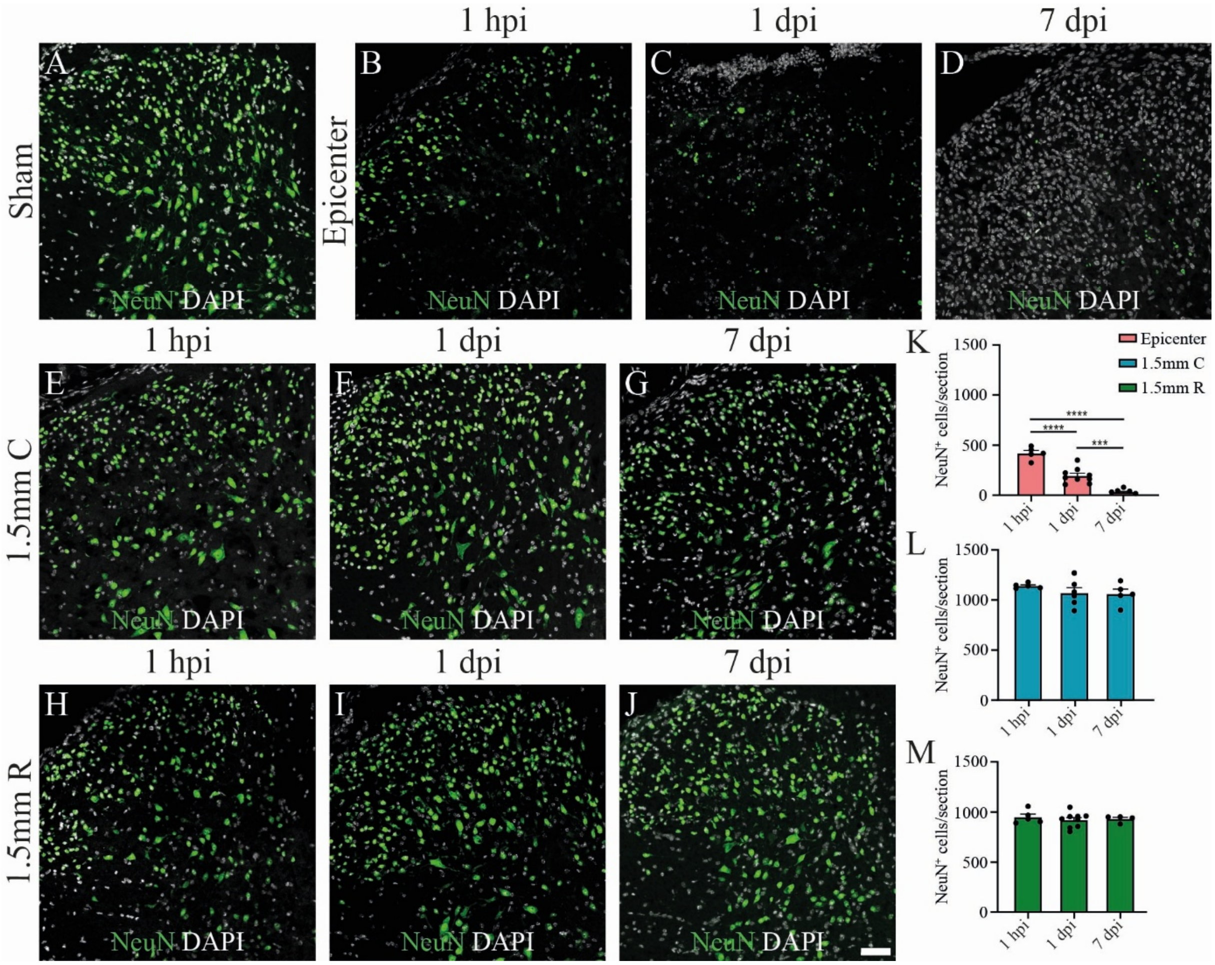


**Supplementary Figure 4. NeuN$^+$ neuronal numbers remain stable at the lesion border despite immunoglobulin internalization after SCI. A**, Representative confocal image of the dorsal horn spinal cord at T9-T10 level of NeuN (in green) and Dapi (in grey). **B-C**, Representative confocal image of the dorsal horn of NeuN (in green) and Dapi (in grey) at the epicenter of the injury at (**B**) 1 hour, (**C**) 1 day and (**D**) 7 days after SCI. **E-G**, Representative confocal image of the dorsal horn of NeuN (in green) and Dapi (in grey) at 1.5mm caudal to the epicenter of the injury at (**E**) 1 hour, (**F**) 1 day and (**G**) 7 days after SCI. **H-I**, Representative confocal image of the dorsal horn of NeuN (in green) and Dapi (in grey) at the epicenter of the injury at (**H**) 1 hour, (**I**) 1 day and (**J**) 7 days after SCI. **K**-**M**, Quantification of NeuN$^+$ cells per section at the epicenter (**K**), 1.5 mm caudal (**L**), and 1.5 mm rostral (**M**). Data represent mean ± SEM. Statistical significance was assessed using one-way ANOVA. Pairwise comparisons and corresponding *p*-values are shown on the graphs, with * $p < 0.05$, ** $p < 0.01$, *** $p < 0.001$, and **** $p < 0.0001$. Scale bars: 50µm in **A-J** (shown in **J**).

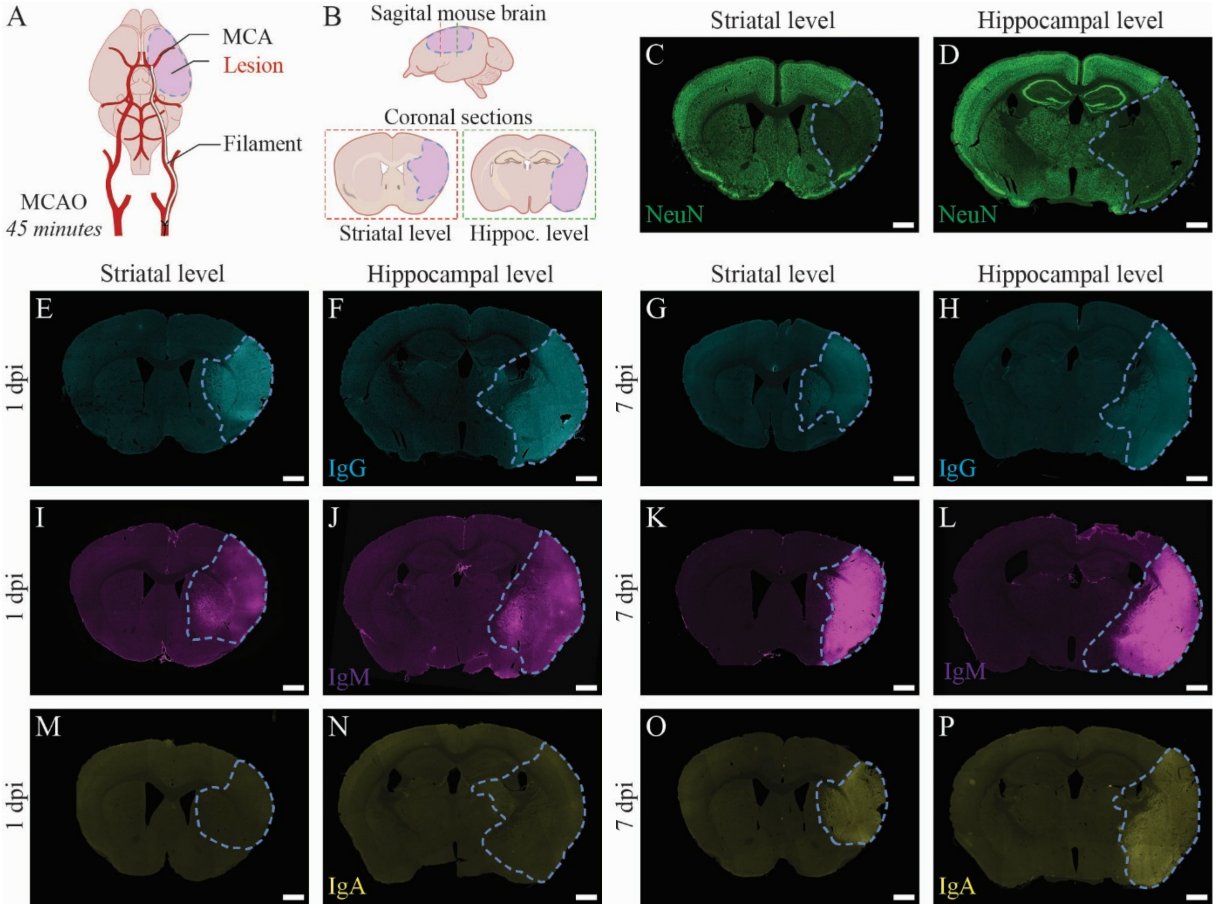


**Supplementary Figure 5. Immunoglobulins accumulate acutely at the ischemic lesion site in the MCAO stroke model. A**, Schematic of the MCAO procedure in mice in which the middle cerebral artery (MCA) was occluded for 45 minutes followed by reperfusion. **B**, Schematic representation of the ischemic lesion after 45 minutes of MCAO, shown in sagittal view (top), and the corresponding coronal sections at the striatal (left) and hippocampal (right) levels where imaging and quantification were performed. **C-D**, Representative confocal images of NeuN immunostaining at the striatal (**C**) and hippocampal (**D**) levels 1 day after MCAO, illustrating how the ischemic lesion site was identified based on decreased NeuN signal. **E-H**, Representative IgG staining (cyan) at the striatal (**E**, **G**) and hippocampal (**F**, **H**) levels at 1 day (**E-F**) and 7 days (**G-H**) after MCAO. **I-L**, Representative IgM staining (magenta) at the same anatomical regions and timepoints. **M-P**, Representative IgA staining (yellow) at the same anatomical regions and timepoints. Scale bars: 500 µm.

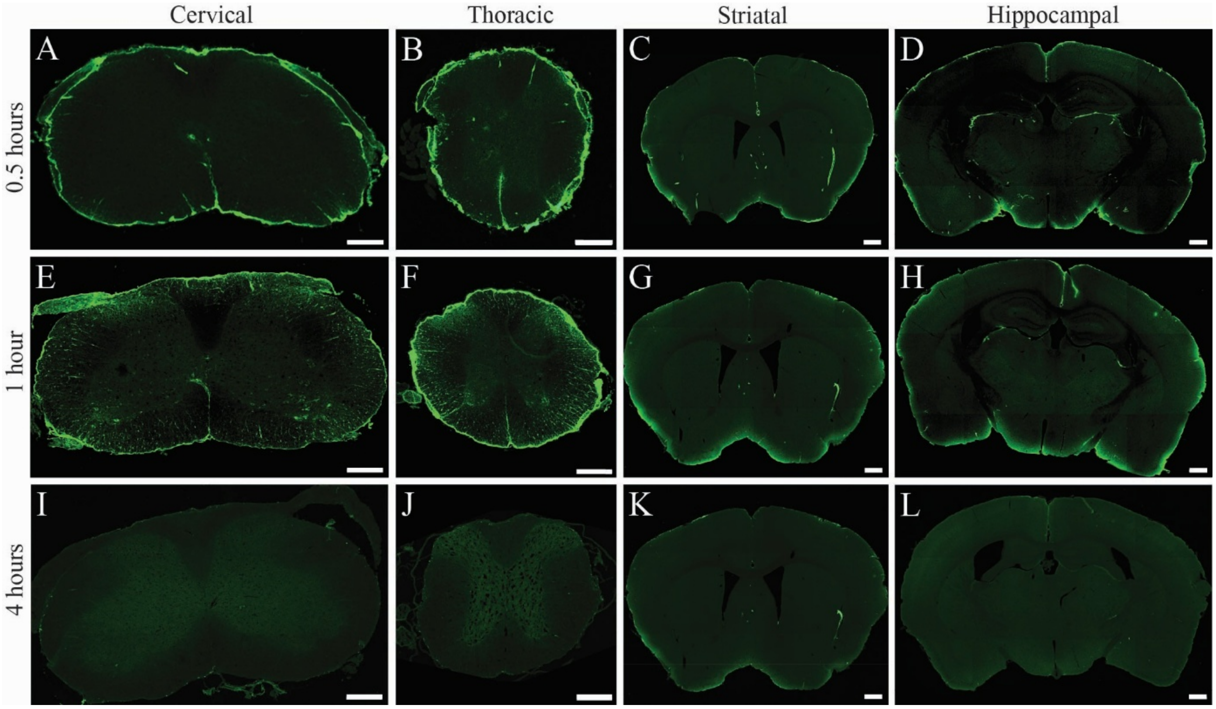


**Supplementary Figure 6. IgG injected into the cisterna magna travels to the brain and spinal cord parenchyma is rapidly cleared from the CNS. A-G**, Representative confocal images showing Alexa488-conjugated IgG in the cervical (**A**, **G**) and thoracic (**B**, **E**, **H**) spinal cord, and in the brain at the striatal (**C**, **F**, **I**) and hippocampal (**D**, **J**) levels at 0.5 hours (**A**-**D**), 1 hour (**E**-**F**), and 4 hours (**G**-**J**) after i.c.m. injection. Scale bars: 250 µm in **A**-**B**, **E**, and **G**-**H**; 500 µm in **C**-**D**, **F**, and **I**-**J**.

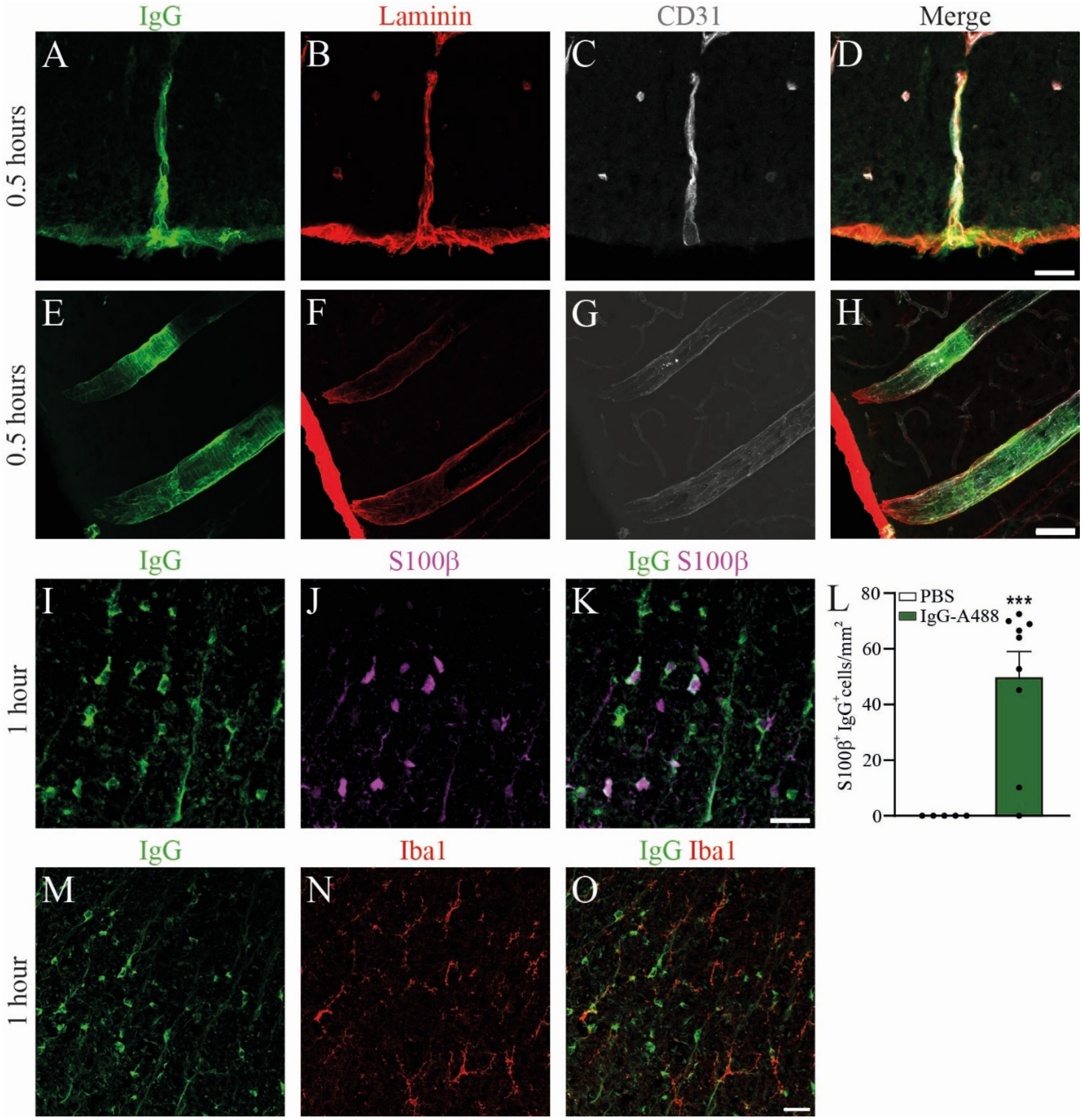


**Supplementary Figure 7. Intracisternally injected IgG reaches the spinal cord and binds to astrocytes and blood vessels in the white matter. A-H**, Representative confocal images of spinal cord (**A-D**) and brain (**E-H**) blood vessels 0.5 hours after i.c.m. injection of Alexa488-conjugated IgG (green). Endothelial cells are labeled with anti-CD31 (grey), and vessel basement membranes with anti-laminin antibody (red). **I-K**, Representative confocal images showing IgG (green) within $S100\beta^{+}$ astrocytes (magenta) in the spinal cord white matter 1 hour after i.c.m. injection of Alexa488-conjugated IgG. **L**, Quantification of $IgG^{+}S100\beta^{+}$ astrocytes in the spinal cord white matter 1 hour post-injection. **M-O**, Representative confocal images showing no colocalization between IgG (green) and $Iba1^{+}$ microglia (red) 1 hour after i.c.m. injection. Data represent mean ± SEM. Statistical significance was assessed using an unpaired t-test with Welch's correction. Pairwise comparisons and corresponding *p*-values are shown on the graphs, with * $p < 0.05$, ** $p < 0.01$, *** $p < 0.001$, and **** $p < 0.0001$. Scale bars: 25 µm in **A-D** (shown in **D**); and 50 µm in **E-H** (shown in **H**); 25µm in **I-K** (shown in **K**) and 25µm in **M-O** (shown in **O**).

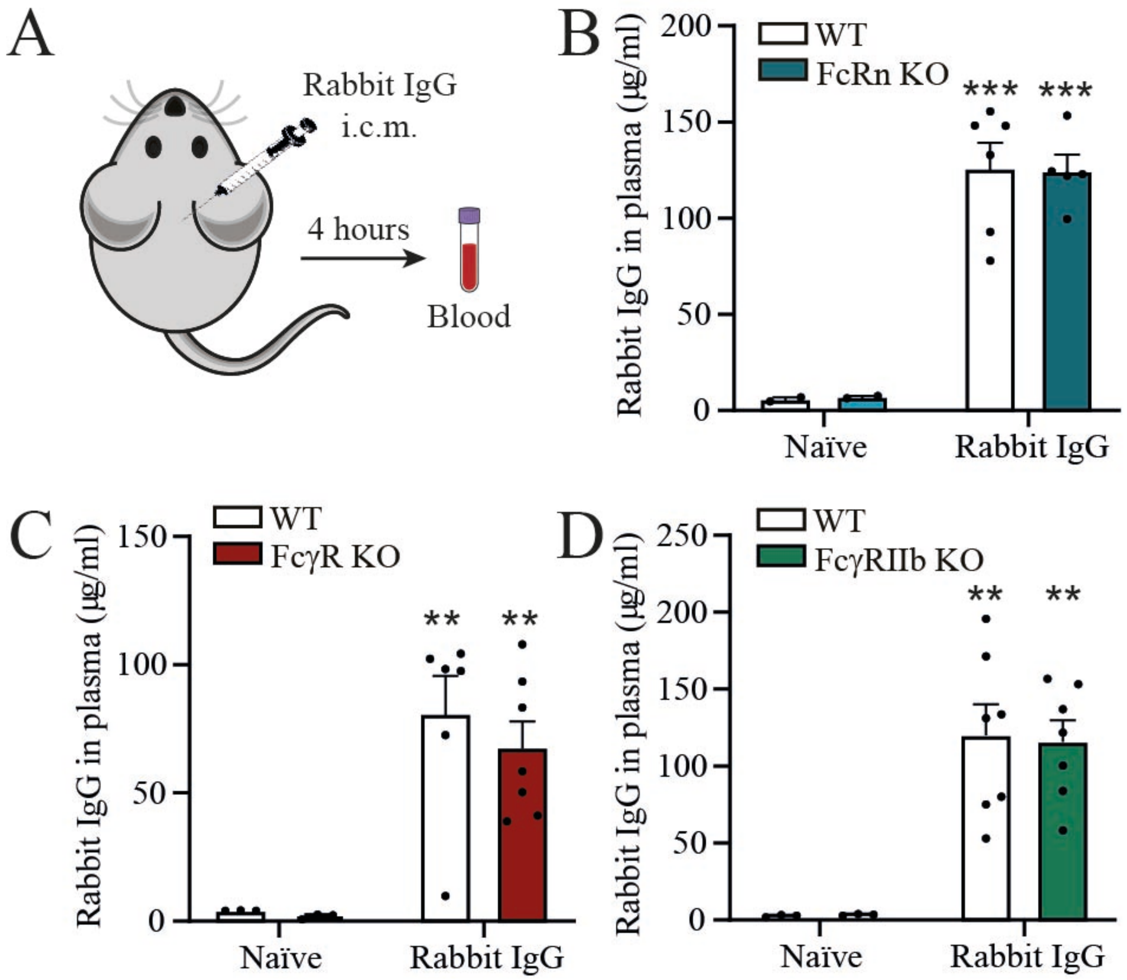


**Supplementary Figure 8. Rabbit IgG injected into the cerebrospinal fluid is rapidly detected in plasma independently of FcRn and Fcγ receptors. A**, Schematic representation of the experimental design. Rabbit IgG was injected intra-cisterna magna (i.c.m.) and blood collected 4 hours later for quantification of rabbit IgG levels in plasma. **B-D**, ELISA quantification of rabbit IgG levels (µg/ml) in the plasma of FcRn-KO (**B**), FcγR-KO (**C**), FcγRIIb-KO (**D**), and their respective wild-type (WT) control mice under naïve conditions or 4 hours after i.c.m. injection of rabbit IgG. Data are shown as mean ± SEM. Statistical analysis was performed using two-way ANOVA with Bonferroni's post-hoc test, with $^{**}p < 0.01$ and $^{***}p < 0.001$ versus the corresponding naïve control group.
